# Single-cell multi-omics defines the cell-type specific impact of splicing aberrations in human hematopoietic clonal outgrowths

**DOI:** 10.1101/2022.06.08.495292

**Authors:** Federico Gaiti, Paulina Chamely, Allegra G. Hawkins, Mariela Cortés-López, Ariel D. Swett, Saravanan Ganesan, Tarek H. Mouhieddine, Xiaoguang Dai, Lloyd Kluegel, Celine Chen, Kiran Batta, John Beaulaurier, Alexander W. Drong, Scott Hickey, Neville Dusaj, Gavriel Mullokandov, Jiayu Su, Ronan Chaligné, Sissel Juul, Eoghan Harrington, David A. Knowles, Daniel H. Wiseman, Irene M. Ghobrial, Justin Taylor, Omar Abdel-Wahab, Dan A. Landau

## Abstract

RNA splicing factors are recurrently affected by alteration-of-function mutations in clonal blood disorders, highlighting the importance of splicing regulation in hematopoiesis. However, our understanding of the impact of dysregulated RNA splicing has been hampered by the inability to distinguish mutant and wildtype cells in primary patient samples, the cell-type complexity of the hematopoietic system, and the sparse and biased coverage of splice junctions by short-read sequencing typically used in single-cell RNA sequencing. To overcome these limitations, we developed GoT-Splice by integrating Genotyping of Transcriptomes (GoT) with enhanced efficiency long-read single-cell transcriptome profiling, as well as proteogenomics (with CITE-seq). This allowed for the simultaneous single-cell profiling of gene expression, cell surface protein markers, somatic mutation status, and RNA splicing. We applied GoT-Splice to bone marrow progenitors from patients with myelodysplastic syndrome (MDS) affected by mutations in the most prevalent mutated RNA splicing factor – the core RNA splicing factor *SF3B1*. High-resolution mapping of *SF3B1*^*mut*^ vs. *SF3B1*^*wt*^ hematopoietic progenitors revealed a fitness advantage of *SF3B1*^*mut*^ cells in the megakaryocytic-erythroid lineage, resulting in an expansion of *SF3B1*^*mut*^ erythroid progenitor (EP) cells. *SF3B1*^*mut*^ EP cells exhibited upregulation of genes involved in regulation of cell cycle and mRNA translation. Long-read single-cell transcriptomes revealed the previously reported increase of aberrant 3’ splicing site usage in *SF3B1*^*mut*^ cells. However, the ability to profile splicing within individual cell populations uncovered distinct cryptic 3’ splice site usage across different progenitor populations, as well as stage-specific aberrant splicing during erythroid maturation. Lastly, as splice factor mutations occur in clonal hematopoiesis (CH) with increased risk of neoplastic transformation, we applied GoT-Splice to CH samples. These data revealed that the erythroid lineage bias, as well as cell-type specific cryptic 3’ splice site usage in *SF3B1*^mut^ cells, precede overt MDS. Collectively, we present an expanded multi-omics single-cell toolkit to define the cell-type specific impact of somatic mutations on RNA splicing, from the earliest phases of clonal outgrowths to overt neoplasia, directly in human samples.

## INTRODUCTION

Genetic diversity in the form of clonal outgrowths has been ubiquitously observed across normal and malignant human tissues^1–13^. Likewise, phenotypic diversity is a hallmark of both normal and malignant tissues in human samples, as has been observed with the widespread application of single-cell RNA sequencing (scRNA-seq)^14–20^. These two axes of cellular diversity likely exhibit complex interplay, as cell state may affect the phenotypic impact of somatic mutations^21^. Recent advances in single-cell multi-omics sequencing have allowed us to reconcile these two aspects of cellular variability in human tissues^15,22,23^, and link genetic variation and transcriptional cell state diversity in somatic evolution. For example, through the application of Genotyping of Transcriptomes (GoT)^15^ technology, which enables genotyping of somatic mutations together with high-throughput droplet-based scRNA-seq, we have previously demonstrated that the effects of somatic mutations on cellular fitness in blood myeloproliferative disorders vary as a function of progenitor cell identity^15^.

Mutations in genes encoding RNA splicing factors serve as an informative example of the challenge of linking genotype to phenotype in complex human tissues. Somatic change-of-function mutations in RNA splicing factors are recurrent in hematologic malignancies^24–26^, highlighting the importance of dysregulated RNA splicing in human hematopoietic disorders. *SF3B1* (splicing factor 3b subunit 1), a core component of the spliceosome complex, is a commonly mutated splicing factor across hematologic malignancies and solid tumors, and is heavily implicated in the pathogenesis of myelodysplastic syndromes (MDS)^27,28^. *SF3B1* mutations also occur in subjects with clonal hematopoiesis (CH), where they confer increased risk of conversion to overt myeloid neoplasms compared to other CH driver mutations^1,2^. *SF3B1* mutations result in incorrect branch point recognition during RNA splicing, often leading to an increased usage of aberrant (or cryptic) intron-proximal 3’ splice sites in hundreds of genes^29^. Such aberrant 3’ splice site recognition typically results in the inclusion of short intronic fragments in spliced mRNA, which most commonly alters the frame of the transcript and renders it a substrate for nonsense mediated mRNA decay (NMD)^30^. Prior work has demonstrated that through mis-splicing, *SF3B1* mutations lead to altered cell metabolism^31^ and disruption of ribosomal biogenesis^32^, leading to the aberrant hematopoietic differentiation typical of MDS. While these are key advances in our understanding of the role of *SF3B1* mutations in MDS development, the mechanisms through which mis-splicing leads to disrupted hematopoietic differentiation in humans remain elusive.

To date, cell culture systems and murine models have been critical for elucidating the role of splicing factor mutations in disordered hematopoiesis. Nonetheless, these methods may not fully recapitulate MDS development in the human context. For example, alternatively spliced genes from murine models of *SF3B1*^*mut*^ MDS, which share some phenotypic similarities with human MDS, show limited overlap with those identified in human samples^33^. The study of splice-altering mutations in humans has been further hampered by three important limitations. First, normal wildtype (WT) and aberrant mutated (MUT) cells are often admixed without discriminating cell surface markers that are required to uniquely isolate MUT cells, limiting the ability to identify signals that can be specifically linked to the *SF3B1*^*mut*^ genotype. This obstacle is magnified in the context of CH where MUT cells commonly constitute a minority of the hematopoietic progenitor population. Second, the hematopoietic differentiation process yields significant complexity of cell progenitor types that further hinders the ability to link mutated genotypes with distinct cellular phenotypes. *SF3B1*^mut^ MDS is indeed associated with a specific clinico-morphological phenotype of refractory anemia and accumulation of ringed sideroblasts^28,34^, strongly suggesting that the interplay between cell identity and *SF3B1* mutations is fundamental in driving disrupted hematopoietic differentiation. Third, scRNA-seq by 3’ or 5’ biased short-read sequencing is limited in its ability to map full-length RNA isoforms and splicing aberrations.

To overcome these limitations and identify cell-identity-dependent mis-splicing mediated by *SF3B1* mutations, we developed GoT-Splice by integrating GoT^15^ with long-read single-cell transcriptome profiling (with Oxford Nanopore Technologies [ONT]) as well as proteogenomics (with CITE-seq)^35^. This allowed for the simultaneous profiling of gene expression, cell surface protein markers, somatic mutation genotyping, and RNA splicing within the same single cell. The application of GoT-Splice to bone marrow progenitor samples from individuals with *SF3B1*-mutated MDS and CH revealed that, while *SF3B1* mutations arise in uncommitted hematopoietic stem progenitor cells (HSPCs), their effect on fitness increases with differentiation into committed erythroid progenitors (EPs), in line with the *SF3B1*^*mut*^-driven dyserythropoiesis phenotype. Importantly, the integration of GoT with full-length isoform mapping via long-read sequencing showed that *SF3B1* mutations exert cell-type specific mis-splicing, already apparent in CH long before disease onset.

## RESULTS

### GoT integrated with proteogenomics reveals increased fitness of *SF3B1*^*mut*^ cells in the erythroid lineage linked to overexpression of cell-cycle and mRNA translation genes

As we have recently demonstrated that the impact of somatic mutations on the transcriptome varies as a function of underlying cell identity in myeloproliferative neoplasms^15^, we hypothesized that an interplay between cell identity and *SF3B1* mutations may drive disrupted hematopoietic differentiation in MDS. To test this, we applied GoT^15^ (**Fig. 1a**) to CD34+ bone marrow progenitor cells from three untreated MDS patients with *SF3B1* K700E mutations (discovery cohort, MDS01-03), as well as a distinct cohort consisting of three MDS patients undergoing treatment (validation cohort, MDS04-06) with erythropoietin (EPO) and/or granulocyte colony-stimulating factor (G-CSF; **Fig. 1b**; **Supplementary Table 1**). As normal hematopoietic development has been extensively studied using flow cytometry cell surface markers, we further integrated GoT with single-cell proteogenomics (CITE-seq^35,36^; **Fig. 1a**). A total of 24,315 cells across the six MDS samples were obtained after sequencing and quality control filtering (**Extended Data Fig. 1a, b**; MDS02 was sequenced in two technical replicates). To chart the differentiation map of the CD34+ progenitor cells, we integrated the data across the primary MDS samples (MDS01-03), as well as the MDS validation samples (MDS04-06), and clustered based on transcriptomic data alone, agnostic to the genotyping and protein information (**Fig. 1c**; **Extended Data Fig. 1c, d**). Using previously annotated RNA identity markers for human CD34+ progenitor cells^37^, validated via Antibody-Derived Tag (ADT) markers in the CITE-seq panel (**Supplementary Table 2, 3**), we identified the expected progenitor subtypes in the primary MDS cohort, along with a population of mature monocytic cells characterized by CD14 expression and lack of CD34 expression often observed in CD34+ sorting of human bone marrow^38^ (**Fig. 1c**; **Extended Data Fig. 2a-c**). Cell clustering was further validated using RNA and ADT multimodal integration (**Extended Data Fig. 2d**). The expected progenitor subtypes were similarly identified in the MDS validation cohort (MDS04-06; **Extended Data Fig. 3a-c**).

**Figure 1.**
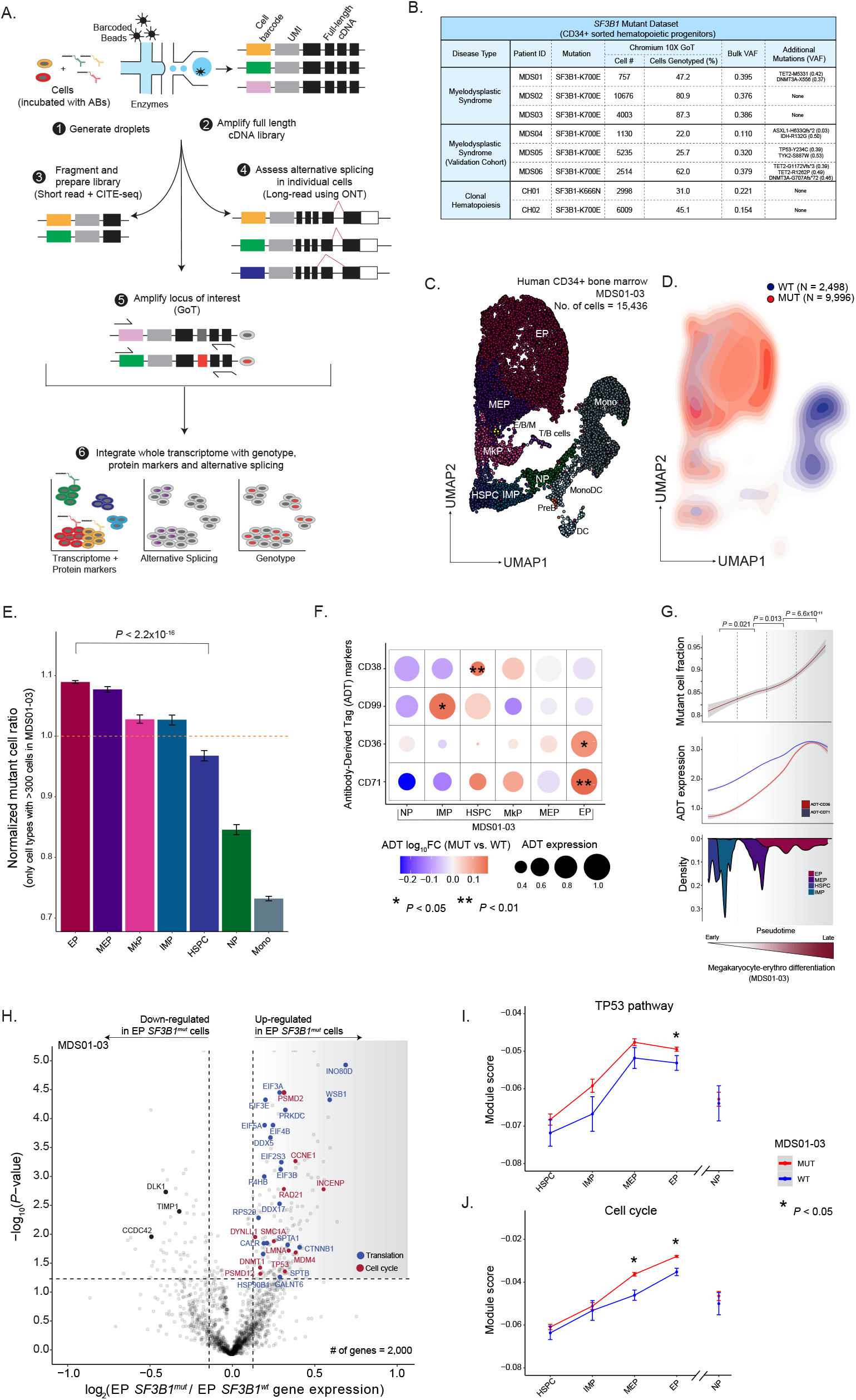
Increased fitness advantage of *SF3B1*^mut^ cells in the megakaryocytic-erythroid lineage. **(A)** Schematic of GoT-Splice workflow. The combination of GoT with CITE-seq and long-read full-length cDNA using Oxford Nanopore Technologies (ONT) enables the simultaneous profiling of protein and gene expression, somatic mutation status, and alternative splicing at single-cell resolution. **(B)** Summary of patient metadata and GoT data (after quality control) for MDS and CH samples with *SF3B1* mutations. **(C)** Uniform manifold approximation and projection (UMAP) of CD34+ cells (*n =* 15,436 cells) from myelodysplastic syndrome patient samples with *SF3B1* K700E mutations (*n =* 3 individuals), overlaid with cluster cell-type assignments. HSPC, hematopoietic stem progenitor cells; IMP, immature myeloid progenitors; MkP, megakaryocytic progenitors; MEP, megakaryocytic-erythroid progenitors; EP, erythroid progenitors; NP, neutrophil progenitors; E/B/M, eosinophil/basophil/mast progenitor cells; T/B cells; Mono, monocyte; DC, dendritic cells; Pre-B, precursors B cells; Mono DC, monocyte/dendritic cell progenitors. **(D)** Density plot of *SF3B1*^*mut*^ vs. *SF3B1*^*wt*^ cells. Genotyping information (MDS01-03) was obtained for 12,494 cells (80.9 % of all cells). **(E)** Normalized frequency of *SF3B1* K700E MUT cells in progenitor subsets with at least 300 genotyped cells. Bars show aggregate analysis of samples MDS01-03 with mean +/-s.e.m. of 100 downsampling iterations to 1 genotyping UMI per cell. Only cell types with >300 cells were used in the analysis. *P*-value from likelihood ratio test of linear mixed model with or without mutation status. **(F)** Differential ADT marker expression between *SF3B1*^*mut*^ and *SF3B1*^*wt*^ cells. Red: higher expression in *SF3B1*^*mut*^ cells; blue: higher expression in *SF3B1*^*wt*^ cells. Size of the dot corresponds to the average expression of ADT marker across cells in a given cell-type. *P*-values determined through permutation testing. **(G)** Mutant cell fraction and ADT expression levels of CD36 and CD71 as a function of pseudotime along the megakaryocyte-erythroid differentiation trajectory for *SF3B1*^*mut*^ and *SF3B1*^*wt*^ cells in MDS01-03. Shading denotes 95% confidence interval. Histogram shows cell density of clusters included in the analysis, ordered by pseudotime. *P*-values were calculated by Wilcoxon rank sum test by comparing mutant cell fraction between pseudotime trajectory quartiles. **(H)** Differential gene expression between *SF3B1*^*mut*^ and *SF3B1*^*wt*^ EP cells in MDS samples. Genes with an absolute log_2_(fold change) > 0.1 and *P*-value < 0.05 were defined as differentially expressed (DE). DE genes belonging to cell cycle (red) and translation (blue) pathways (Reactome) are highlighted (BH-FDR < 0.2). **(I)** Expression (mean +/-s.e.m.) of TP53 pathway related genes (Reactome) between *SF3B1*^*mut*^ and *SF3B1*^*wt*^ cells in progenitor cells from MDS01-03 samples. Red: module score in *SF3B1*^*mut*^ cells; blue: module score in *SF3B1*^*wt*^ cells. *P*-values from likelihood ratio test of linear mixed model with or without mutation status. **(J)** Same as **(I)** for expression of cell cycle related genes (Reactome) between *SF3B1*^*mut*^ and *SF3B1*^*wt*^ cells in progenitor cells from MDS01-03 samples.

Genotyping data were available for 15,650 MDS cells (64.4% across MDS01-06) through GoT (**Fig. 1b**; **Extended Data Fig. 4a-d**). The per-patient mutant cell fractions obtained through GoT were highly correlated with the variant allele frequencies (VAFs) obtained through bulk sequencing of matched unsorted peripheral blood mononuclear cells (Pearson’s r = 0.81, *P*-value = 0.008; **Extended Data Fig. 4a**). Projection of the genotyping information onto the differentiation map demonstrated MUT and WT cells co-mingled throughout the differentiation topology (**Extended Data Fig. 4c, d**), highlighting the need for single-cell multi-omics to link genotypes with cellular phenotypes in *SF3B1*^*mut*^ MDS. Although MUT cells were found across CD34+ progenitor cells, we observed an accumulation of MUT cells along the erythroid trajectory (**Fig. 1d**), suggesting that *SF3B1* mutant cell frequency (MCF) varies as a function of the progenitor subtype. To confirm this, we evaluated the MCF across the different prevalent progenitor cell types (limited to progenitor subsets with > 300 cells). Of note, as cells may display variable expression of *SF3B1* itself, we performed amplicon UMI-downsampling to exclude sampling biases given the heterozygosity of the mutated allele as a potential confounder for observed differences in MCF (see Methods). Across samples, we observed a significant increase in MCF in the megakaryocyte-erythroid lineage with the highest MCF observed in EPs compared to HSPCs (*P-*value *<* 10^−16^; **Fig. 1e**; **Extended Data Fig. 4e**), consistent with the erythroid lineage-specific impact of mutated *SF3B1*^39,40^.

The ability to layer protein measurements on top of GoT data further allowed us to identify differentially expressed proteins between MUT and WT cells within each progenitor subset. After quality control filtering for ADT markers with adequate expression in at least two major progenitor subtypes (see Methods), protein expression was highest in the expected cell types, and correlated with mRNA expression, both at the individual cell as well as cell-type level, comparable to previous data^35^ (**Extended Data Fig. 5a, b**). We directly compared protein expression between MUT and WT cells, accounting for sample-to-sample variability in mutated cells through downsampling (see Methods), and observed differential expression of CD38, CD99, CD36 and CD71 in at least one progenitor cell-type (**Fig. 1f**; **Supplementary Table 4**). CD38 is a known marker for the transition of primitive CD34+ stem and progenitor cells into more committed precursor cells^37,41,42^. Its overexpression in *SF3B1*^*mut*^ is consistent with the observed higher MCF in committed progenitor subsets. CD99, over-expressed in MUT immature myeloid progenitor cells (IMP) cells, was previously noted to be overexpressed in both AML and MDS stem cells, serving as a potential therapeutic target of malignant stem cells^43,44^. Finally, CD36 and CD71, erythroid lineage markers, were found to be over-expressed in MUT EPs when compared to WT EPs, consistent with the *SF3B1*^*mut*^-driven dyserythropoiesis phenotype. We further leveraged these erythroid maturation cell surface protein markers to validate pseudo-temporal (pseudotime) ordering of the continuous process of erythroid maturation^45^ (**Extended Data Fig. 5c**). This analysis revealed an increase in MCF along erythroid lineage maturation (**Fig. 1g**), confirming that *SF3B1* mutational fitness increases with differentiation into committed EPs.

To further explore *SF3B1* driven transcriptional dysregulation in committed EPs, we performed differential gene expression analysis between *SF3B1*^*mut*^ and *SF3B1*^*wt*^ cells. Mutated EPs upregulated genes encoding important translation and ribosome biogenesis factors (FDR < 0.2; **Fig. 1h**; **Supplementary Table 5, 6**), including a number of eukaryotic initiation factors (*e*.*g*., *EIF3, EIF5*), DEAD-box helicases (*e*.*g*., *DDX5, DDX17*), and ribosome subunits (*e*.*g*., *RPS29*). This dysregulation of translational activity, or ribosomal stress signal, is evocative of studies showing that translational regulation is critical during hematopoiesis^46–49^, and may lead to cell- and tissue type–restricted activation of *TP53* signaling pathway in myeloid disease^50–55^. Specifically, cells that require high levels of protein synthesis, such as erythroid progenitors, may be more sensitive to changes caused by translational dysfunction^56^. In line with this notion, *TP53* gene target upregulation in *SF3B1*^*mut*^ cells was more prominent in the megakaryocyte-erythroid lineage, with no increased expression of *TP53*-related genes in earlier progenitors (HSPCs) or in neutrophil progenitors (NPs) compared to WT cells (**Fig. 1i**). Our results therefore establish a molecular phenotype for the *SF3B1* mutation in human bone marrow progenitors, potentially phenocopying translational dysregulation typically observed in ribosomopathies driven by germline deleterious mutations of ribosomal subunits^56^.

Mutated EPs also upregulated genes involved in cell-cycle and checkpoint control (FDR < 0.2; **Fig. 1h, j**; **Supplementary Table 5, 6**). In particular, we observed an increase in expression of *CCNE1*, a positive regulator of the G1/S transition of the cell cycle^57^, and *MDM4*. The latter gene works together with *TP53* during the G1/S checkpoint of the cell cycle to determine cell fate by regulating pathways such as DNA repair, apoptosis, and senescence^58^. Increased expression of *MDM4* during ribosomal stress^59^ prevents TP53 degradation and blocks subsequential inactivation of p21, resulting in a sustained cell proliferation^60^. Together, the combined upregulation of *TP53, CCNE1*, and *MDM4* in mutated EPs may therefore lead to cell survival and accumulation rather than cell death, supporting the finding that *SF3B1* mutations impart a greater fitness advantage specifically in the erythroid lineage.

### GoT-Splice links somatic mutations, alternative splicing, and cellular phenotype at single-cell resolution

The integration of GoT with single-cell proteogenomics data revealed that *SF3B1* mutations reshape hematopoietic differentiation and mediate cell-identity-dependent transcriptional changes (**Fig. 1**). Given the pivotal role of *SF3B1* in mRNA splicing, we next explored how mis-splicing may serve as a link between genotypes and cellular phenotypes. Indeed, *SF3B1* mutations promote recognition of alternative branch points, most commonly leading to increased usage of aberrant 3’ splice sites^29^. However, previous studies in primary human samples have been performed on bulk samples admixing MUT and WT cells as well as progenitor subtypes^30,32,61,62^. Conversely, short-read sequencing typically employed in scRNA-seq does not adequately cover splice junctions. Recent advances suggest that long-read integration into scRNA-seq may overcome these limitations^63–67^. We therefore integrated GoT with full-length ONT long-read sequencing, allowing for high-throughput, single-cell integration of genotype, cell surface proteome, gene expression, and mRNA splicing information (GoT-Splice; **Fig. 1a**). We note that single-cell cDNA sequencing with ONT presents unique challenges, as cDNA amplification artifacts are still productively sequenced when using standard ONT ligation chemistry. This leads to a high fraction of uninformative reads in the highly amplified single-cell libraries. To enhance ONT efficiency, we incorporated a biotin enrichment step using on-bead PCR to selectively amplify full-length reads containing intact cell barcodes and unique molecular identifiers^64^ (UMIs; **Fig. 2a**). This approach increased the yield of full-length reads from 50.4 +/-2.7 to 77.6 +/-2.0 (mean +/-s.e.m.) percent of all sequenced reads. Thus, GoT-Splice delivers high-resolution single-cell full-length transcriptional profiles that are comparable with short-read sequencing (**Fig. 2b, c**). To accurately identify splice junctions using single-cell long-read sequencing, we developed an analytical pipeline that leverages the recently published SiCeLoRe pipeline^64^ (**Extended Data Fig. 6a**). To reduce alignment noise, we generated a splice junction reference identified in single-cell SMART-seq2 data from human CD34+ cells with no *SF3B1* mutation (see Methods). Next, we carried out intron-centric junction calling which allows for the independent measurement of splicing at both the 5’ and 3’ ends of each intron. This allows for an unbiased assessment of junctions and a greater accuracy in measuring the degree of mis-splicing of a particular transcript when compared to exon-centric quantification approaches^68^, which are typically used for cassette exon usage profiling and rely on predefined transcript models or splicing events, both of which may be inaccurate or incomplete^69,70^. As anticipated, we observed a 4-fold increase in the number of junctions per cell detected using full-length long-read sequencing over short-read, despite lower absolute number of UMI/cell (**Fig. 2d**). Additionally, GoT-Splice afforded greater coverage uniformity across the entire transcript, compared to 3’-biased coverage in short-read sequencing (**Fig. 2e**).

**Figure 2.**
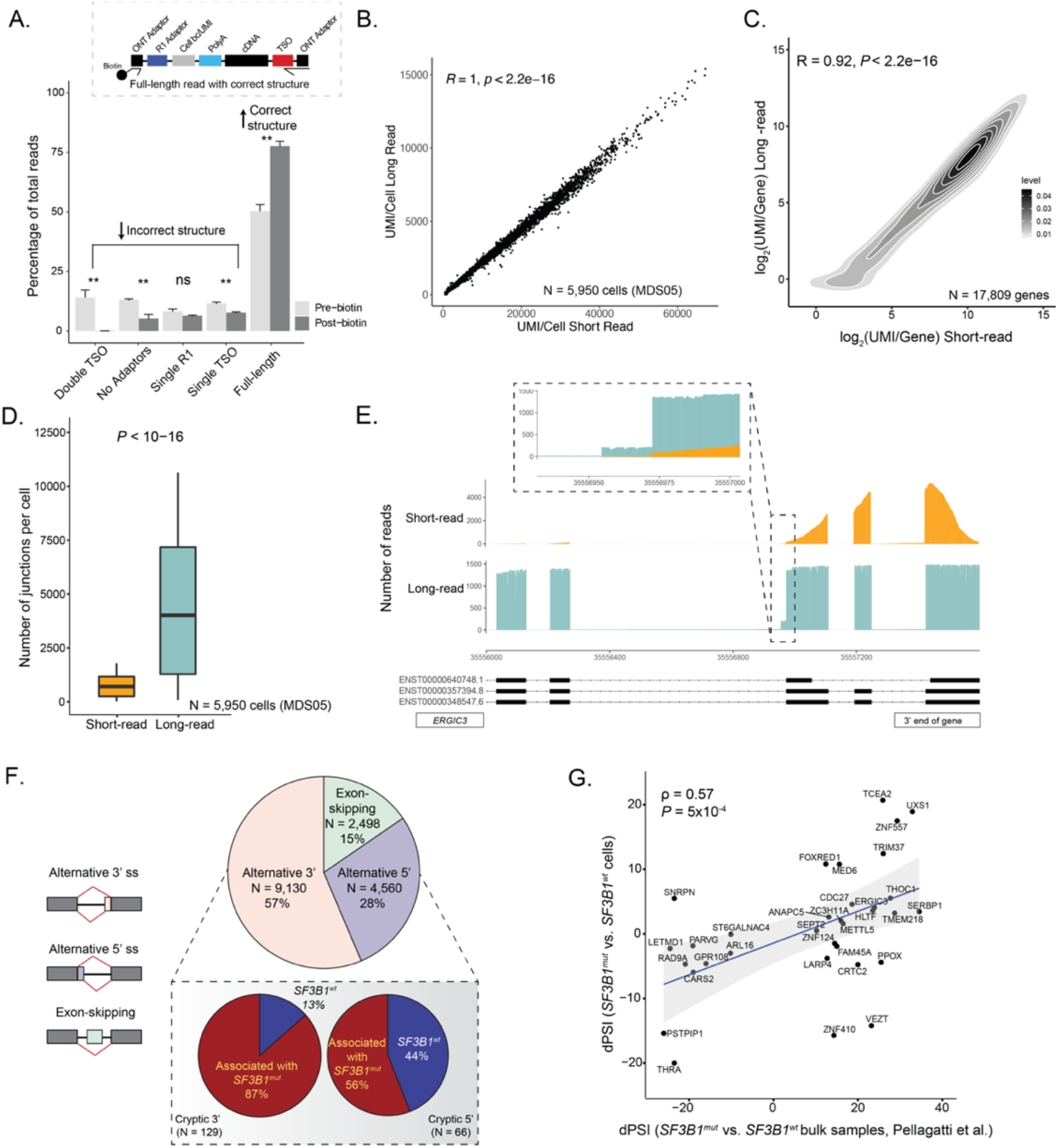
Simultaneous profiling of gene expression, cell surface protein markers, somatic mutation status and alternative splicing at single-cell resolution. **(A)** A comparison of the percentage of ONT reads with either incorrect structure (double TSO, no adaptors, single R1 or single TSO) or correct structure (full-length reads) both before and after the inclusion of a biotin enrichment protocol step during preparation for sequencing. Bars show the aggregate analysis of *n* = 5 samples with mean +/-s.d. of the percentage for each category. **(B)** Scatter plot of the correlation between the number of UMIs/cell detected in long-read ONT vs. short-read Illumina data for cells that were sequenced across both platforms for sample MDS05. **(C)** Density plot of the correlation between the number of UMIs/gene detected in long-read ONT vs. short-read Illumina data for sample MDS05. **(D)** Number of splice junctions captured in the full-length long-read ONT data compared to short-read sequencing data, showing that GoT-Splice allows for a significant increase in the number of junctions captured per cell. **(E)** GoT-Splice provides greater sequencing coverage uniformity compared to inadequate coverage of short-read sequencing over splice junctions, as exemplified here for the *ERGIC3* gene. **(F)** Pie chart summarizing the distribution of different alternative splicing events detected after junction annotation. *Inset*: Pie chart summarizing the differences in the usage of cryptic 3’ and 5’ splice site events between *SF3B1*^*mut*^ and *SF3B1*^*wt*^ cells measured with a dPSI (*SF3B1*^*mut*^ PSI - *SF3B1*^*wt*^ PSI). Associated with *SF3B1*^*mut*^: +ve dPSI; associated with *SF3B1*^*wt*^: -ve dPSI. **(G)** Comparison of delta percent spliced-in (dPSI) values of shared cryptic 3’ splicing events identified in the MUT vs. WT cell comparison from GoT-Splice of *SF3B1*^*mut*^ MDS01-03 samples and in the *SF3B1*^*mut*^ vs. *SF3B1*^*wt*^ bulk comparison from bulk RNA-sequencing of CD34+ cells of MDS samples in Pellagatti et al.^32^. Correlation coefficient *ρ* calculated using Spearman’s correlation and *P*-value derived from Student’s t-distribution.

The most common mis-splicing events observed in MDS *SF3B1*^*mut*^ cells involved alternative 3’ splice sites, accounting for 57% of alternative splicing events (**Fig. 2f**), consistent with prior reports^29,71^. Notably, the usage of such alternative 3’ splice sites was not observed in a CD34+ sample with no *SF3B1* mutation (**Extended Data Fig. 6b**). ONT long-read sequencing also allowed us to quantify the presence of different splicing events across the same mRNA transcript. While only one aberrant 3’ splice site event was observed for the majority of mRNA transcripts, we identified a total of 428 genes (21.4% of the total number of genes with at least one cryptic 3’ splice site) with more than one aberrant 3’ splice site event. Interestingly, these cryptic 3’ splicing events tend to appear in different copies of the transcript (**Extended Data Fig. 6c**), highlighting the unique advantages of long-read sequencing in this context. Consistent with previous MDS bulk sequencing data^72,73^, we observed a relative enrichment of purines upstream of the aberrant 3’ splice site when compared to the canonical 3’ splice site (**Extended Data Fig. 6d**).

We next leveraged the unique ability of GoT-Splice to resolve differential splice junction usage between *SF3B1*^*mut*^ and *SF3B1*^*wt*^ cells within the same primary human sample (see Methods). Of the differentially mis-spliced cryptic 3’ splice sites (those 0-100bp from the canonical splice site) between MUT and WT cells, 87% were used more highly in *SF3B1*^*mut*^ cells (**Fig. 2f, *inset***), aligning with known characteristics of *SF3B1* mutations. Furthermore, we observed a high correlation between GoT-Splice delta PSI (dPSI; percent spliced in) measurements obtained by comparing *SF3B1*^*mut*^ and *SF3B1*^*wt*^ cells, and dPSI derived from bulk RNA-sequencing of CD34^+^ cells from *SF3B1*^*mut*^ vs. *SF3B1*^*wt*^ MDS samples^32^ for shared cryptic 3’ splice sites (**Fig. 2g**). In line with previous work, the majority of these cryptic 3’ splice sites were found to be ∼15-20 bps upstream of the canonical 3’ site^29^ (**Fig. 3a; Extended Data Fig. 7a-d**). GoT-Splice enabled the visualization of cryptic 3’ splice sites in *SF3B1*^*mut*^ vs. *SF3B1*^*wt*^ cells, highlighting the striking increased usage of cryptic 3’ splice sites specific to *SF3B1*^*mut*^ (**Fig. 3b**). Altogether, GoT-Splice extends the ability to connect somatic mutations not only to transcriptional and cell surface protein marker phenotypes, but also to single-cell mapping of splicing changes.

**Figure 3.**
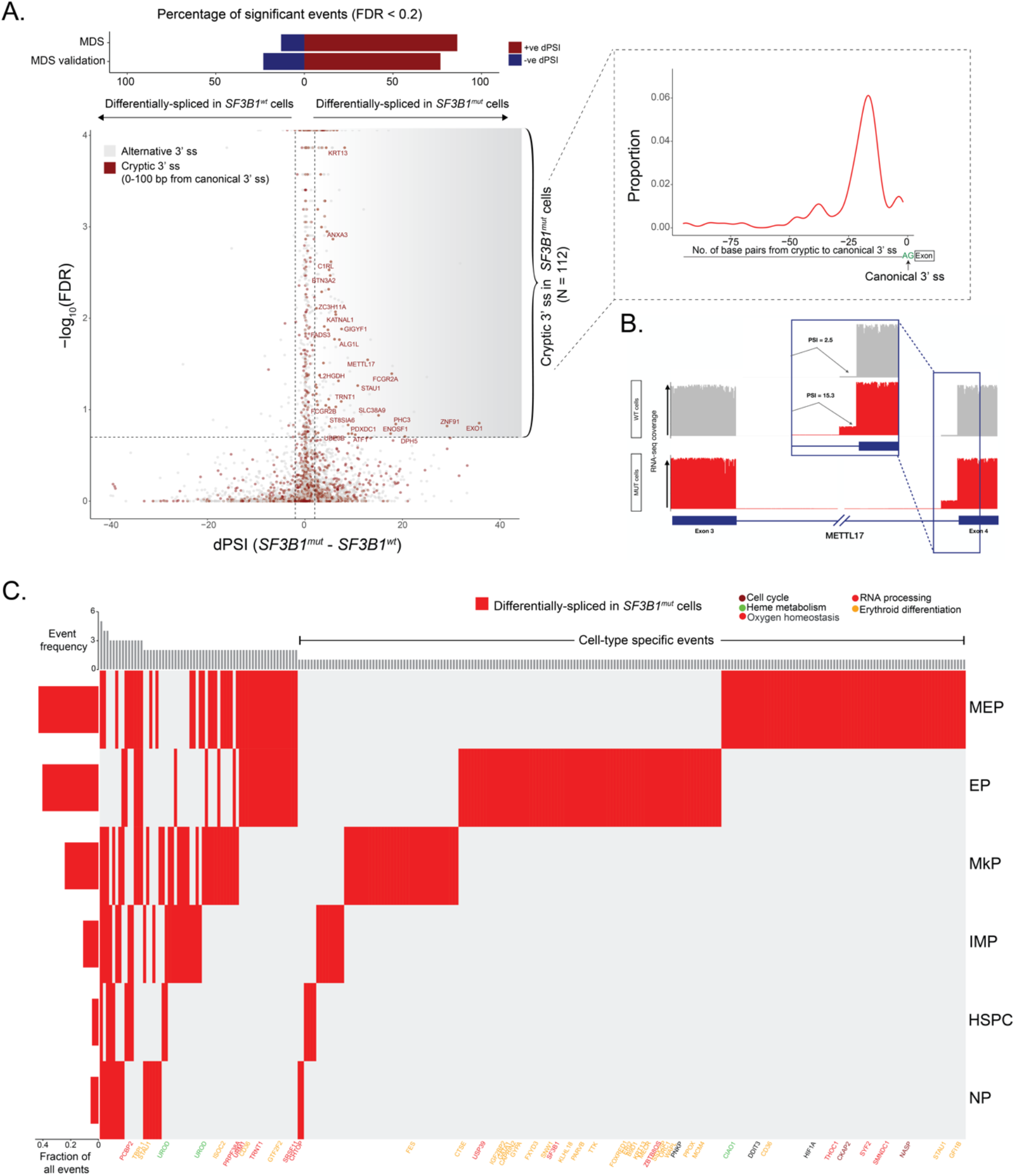
Progenitor cell-type specific mis-splicing in *SF3B1*^*mut*^ MDS. **(A)** Differential splicing analysis between *SF3B1*^*mut*^ and *SF3B1*^*wt*^ cells across MDS samples. Junctions with an absolute dPSI > 2 and BH-FDR adjusted *P*-value < 0.2 were defined as differentially spliced. *Top:* Bars showing the percentage of genes differentially spliced in *SF3B1*^*mut*^ and *SF3B1*^*wt*^ cells in the MDS and MDS validation cohorts. *Inset*: Expected peak in the number of identified cryptic 3’ splice sites at the anticipated distance (15-20 base pairs) upstream of the canonical 3’ splice site in *SF3B1*^*mut*^ cells. **(B)** Sashimi Plot of *METTL17* intron junction with an *SF3B1*^*mut*^ associated cryptic 3’ splice site showing RNA-seq coverage in *SF3B1*^*mut*^ vs. *SF3B1*^*wt*^ cells within MDS samples. *Inset:* Expected marked increase in the PSI value for the usage of this cryptic 3’ splice site in *SF3B1*^*mut*^ cells. **(C)** Representation of dPSI values between *SF3B1*^*mut*^ and *SF3B1*^*wt*^ cells for cryptic 3’ splicing events identified in the main progenitor subsets across MDS samples. Rows correspond to cryptic 3’ junctions found to be differentially spliced in at least one cell-type, with *P*-value <= 0.05 and dPSI >= 2. Columns correspond to cell-type. Genes that belong to pathways cell cycle (purple), heme metabolism (green), oxygen homeostasis (black), RNA processing (red) and erythroid differentiation (yellow) are highlighted. The left bar plots show the fraction of differentially spliced cryptic 3’ splice sites per cell. Top bar plots quantify the total number of cell types where an event is differentially spliced, with the cell-type specific events located to the right side of the plot.

### GoT-Splice shows progenitor-specific patterns in *SF3B1*^*mut*^-mis-splicing

An important advantage of GoT-Splice is the ability to detect splicing changes at single-cell resolution, which enables the comparison of alternative splicing aberrations between MUT and WT cells within specific cell subsets (**Fig. 3c, Supplementary Table 7**). We identified both shared and unique *SF3B1*^*mut*^ cryptic 3’ splice site events across progenitor subtypes. The usage of cryptic 3’ splice sites was highest along the megakaryocyte-erythroid lineage, with *SF3B1*^*mut*^ MEPs and EPs accounting for the majority of cell-type specific cryptic 3’ splice site events, highlighting the specific impact of *SF3B1* mutations on the erythroid lineage. These progenitor specific patterns in *SF3B1*^*mut*^ mis-splicing were further detected in the validation cohort of MDS patient samples (MDS04-06; **Extended Data Fig. 7e, f**). In both MDS cohorts, progenitor specific cryptic 3’ splice sites involved genes related to cell cycle (*e*.*g*., *CENPT*)^74^, RNA processing (*e*.*g*., *CHTOP, SF3B1*^75^, *SRSF11, PRPF38A*), erythroid differentiation (*e*.*g*., *CD36, FOXRED1, GATA1*^34,76,77^), and heme metabolism (*e*.*g*., *UROD, PPOX, CIAO1*) (**Fig. 3c**; **Extended Data Fig. 7e, f**; **Supplementary Table 7, 8**). Many of these genes and pathways have previously been reported to be disrupted by alternative splicing in bulk studies of *SF3B1*^*mut*^ MDS samples^32^, but their cell-type specificity was unknown. For instance, while the alternative splicing event in *SF3B1* itself has been suggested before as being neoplasm-specific, here we narrowed down its erythroid-specific pattern. This isoform – *SF3B1ins –* is predicted to affect splicing by impairing U2 snRNP assembly^75^, likely contributing to the enhanced mis-splicing dysregulation in the megakaryocyte-erythroid lineage. In addition, cell cycle plays a critical role in the terminal differentiation of hematopoietic stem cells^78^ and RNA processing, erythroid differentiation, and heme metabolism pathways are directly linked to the regulation of erythropoiesis^79–81^. To further validate cell-type specificity of mis-splicing events, we compared the genes with cryptic 3’ splice site events unique to MEPs and EPs in the two distinct MDS cohorts and observed significant overlap of megakaryocyte-erythroid lineage-specific aberrantly spliced genes between the discovery and the validation MDS cohorts (*P*-value = 0.00029, Fisher’s exact test, with 46.8% of the cryptically spliced genes in MDS also aberrantly spliced in the MDS validation cohort). In contrast, no significant overlap was observed when comparing the genes with cryptic 3’ splice site events unique to MEPs and EPs in the MDS discovery cohort to genes with cryptic 3’ splice sites unique to earlier progenitor cells in the MDS validation cohort (1.6% overlap; *P*-value = 0.46, Fisher’s exact test; **Extended Data Fig. 7f**). These findings reveal that alternative splicing is cell-type and differentiation-stage dependent^27,82–84^.

Of note, erythropoiesis occupies a continuum of cell states and is dependent on a series of transcriptional changes that occur along a continuous trajectory^45^. Analyzing the *SF3B1*^*mut*^ mis-splicing along this continuum (**Fig. 4a**) revealed that some erythroid differentiation and heme metabolism genes can be mis-spliced more frequently at the earliest stages of EP maturation (*e*.*g*., *UROD* and *FOXRED1*^85^), while others display increased mis-splicing in the more differentiated EPs (*e*.*g*., *GYPA* and *PPOX*). *UROD* is part of the heme biosynthesis pathway and not only is heme an important structural component of erythroid cells but it also plays a regulatory role in the differentiation of erythroid precursors^86^. *PPOX* encodes for an enzyme involved in mitochondrial heme biosynthesis and, as such, its degradation leads to ineffective erythropoiesis and accumulation of iron in the mitochondria typical of MDS with ring sideroblast clinical phenotype^87^. These results provide evidence that disruptive and pathogenic *SF3B1*^*mut*^-driven mis-splicing impacts key mediators of hemoglobin synthesis and erythroid differentiation at all stages of erythroid maturation^88,89^.

**Figure 4.**
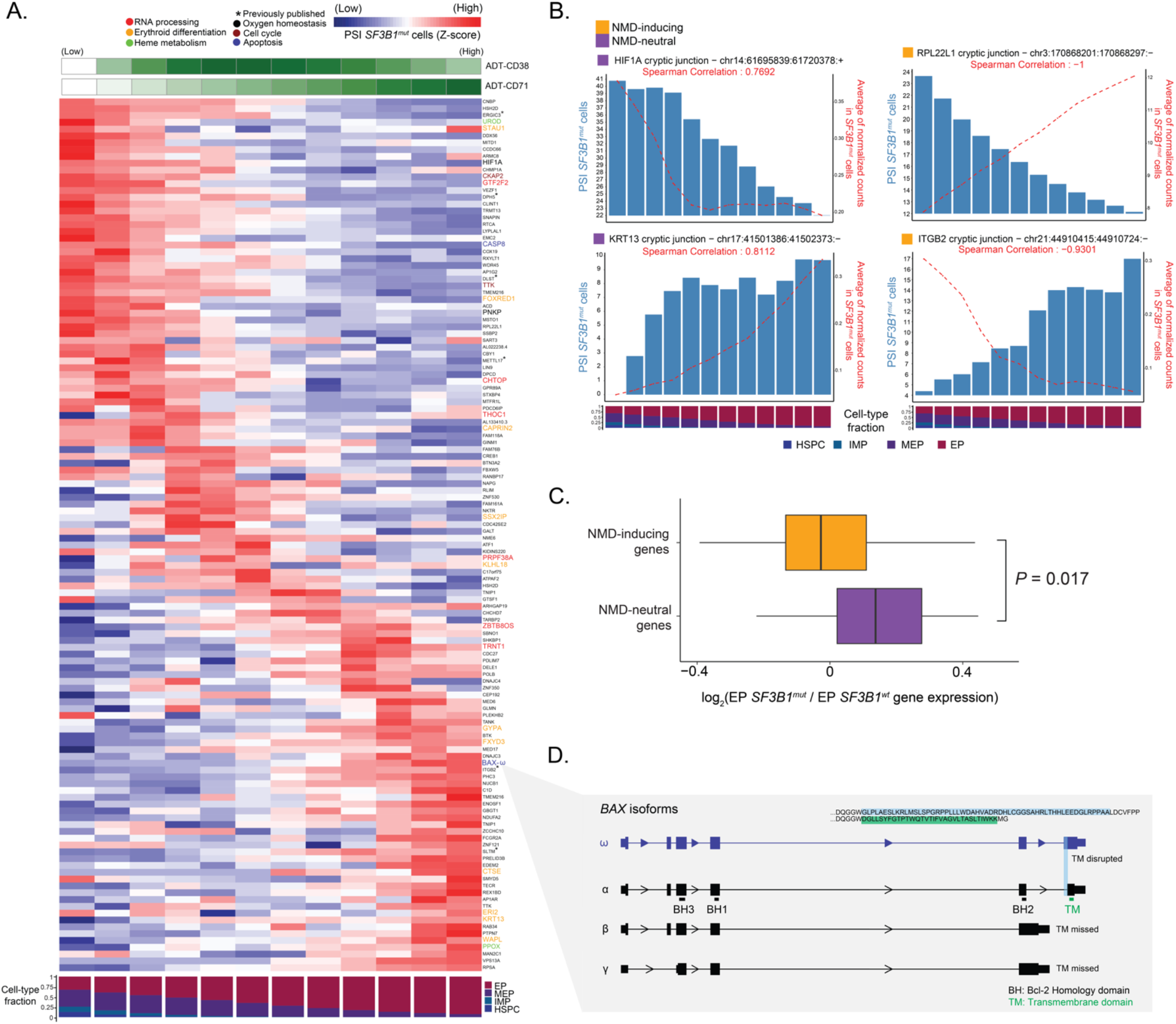
*SF3B1*^*mut*^-associated mis-splicing changes along the continuum of erythropoiesis. **(A)** Percent spliced-in (PSI) of junctions in *SF3B1*^*mu*t^ cells along the hematopoietic differentiation trajectory (HSPCs, IMPs, MEPs, EPs). Rows (z-score normalized) correspond to cryptic 3’ splice sites; columns represent the PSI for the usage of a given cryptic 3’ splice site in each window (size of 3000 *SF3B1*^*mut*^ cells, sliding by 300 *SF3B1*^*mut*^ cells). Only junctions found to be differentially spliced in at least one cell-type with a dPSI > 2 were used in the analysis. The ADT expression of erythroid lineage marker CD71, along with the fraction of cell types in each window, is shown. Rows are ordered according to the peak in PSI. Genes that belong to pathways cell cycle (purple), heme metabolism (green), oxygen homeostasis (black), RNA processing (red), erythroid differentiation (yellow) and apoptosis (blue) are highlighted. **(B)** Examples of mis-spliced genes at different stages of erythroid maturation. Bars represent PSI in *SF3B1*^*mut*^ cells. Red lines represent ONT expression of the given junction in *SF3B1*^*mut*^ cells. **(C)** Fold change (log_2_) of gene expression between *SF3B1*^*mut*^ and *SF3B1*^*wt*^ EP cells in NMD-inducing vs. NMD-neutral genes. **(D)** Gene model of *BAX* and relevant isoforms. Characteristic domains and their location are highlighted in *BAX-α*, the main isoform. The cryptic 3’ splicing event on the terminal exon defines the *BAX-ω* isoform, characterized by the disruption of the transmembrane domain (TM) as a result of a frameshift.

We further noted that the degree of mis-splicing of a particular transcript (measured via PSI) positively correlated with its expression across the erythroid differentiation trajectory in some cases. In others, mis-splicing was anti-correlated with gene expression, often in cryptic 3’ splice site events that are predicted to lead to transcript degradation by the NMD pathway (**Fig. 4b** for representative examples). Cryptic 3’ splice sites result in the inclusion of short intronic fragments in mRNA and often introduce a premature termination codon (PTC)^90–92^. mRNAs harboring an NMD-inducing PTC located ≥50 bps upstream of the last exon–exon junction are predicted to undergo NMD, which in turn prevents the production of potentially aberrant proteins. In contrast, mRNAs harboring an NMD-neutral PTC, which is generally located ≤50 bps upstream of the last exon–exon junction or in the last exon, fail to trigger NMD and produce dysfunctional proteins^93,94^. We classified cryptic 3’ splice sites detected in the MDS samples into three major groups: (i) NMD-inducing event (due to the introduction of a PTC); (ii) NMD-neutral with a frameshift event; and (iii) NMD-neutral with no frameshift event (**Supplementary Table 7**). In accordance with previous reports^71^, of the 421 cryptic 3’ splice sites significantly associated with the *SF3B1*^*mut*^ cells, 228 (54%) of these were classified as NMD-inducing events while the remaining 193 (46%) were NMD-neutral (60 events involving a frameshift and 133 events were in-frame). As expected, we observed a significant decrease in the expression of genes harboring NMD-inducing events compared with those harboring NMD-neutral events (*P*-value = 0.017, Mann Whitney U test; **Fig. 4c**).

NMD-inducing events affected genes including *UROD, GYPA, FOXRED1* and *PPOX* – key genes in erythroid development. The loss of these transcripts via NMD^95,96^ may thus contribute to disrupted terminal differentiation of EPs. Notable among NMD-neutral affected genes, we identified *BAX*, a member of the Bcl-2 gene family and transcriptional target of *TP53. BAX* is a vital component of the apoptotic cascade and in turn plays an important role in balancing the control of survival, differentiation and proliferation of EPs at later stages of erythropoiesis^97^ (**Fig. 4a**). The identified *BAX* cryptic 3’ splice site, though NMD-neutral, causes a frameshift in the last exon, disrupting the C-terminus of the protein. This *BAX* isoform, previously denoted as *BAX-ω* (**Fig. 4d**), has been shown to protect cells from apoptotic cell death^98,99^. Interestingly, a recent study revealed C-terminal *BAX* mutations in myeloid clones that arise in chronic lymphocytic leukemia patients upon prolonged exposure to venetoclax, demonstrating a role for BAX c-terminal alterations in conferring a survival advantage to myeloid cells with this pro-apoptotic treatment. Of note, early clinical observations reported lower response to venetoclax in *SF3B1*^*mut*^ AML^100,101^, consistent with a potential anti-apoptotic effect of *BAX-ω*. Together, these findings suggest a potential mechanism underlying the erythroid-dysplasia phenotype of *SF3B1*^*mut*^ MDS. Despite the injury to translational machinery (**Fig. 1h-i**), *SF3B1*^*mut*^ EPs may gain some degree of protection against cell death due to the presence of isoform *BAX-ω*, arising from aberrant splicing.

### Accumulation of *SF3B1*^*mut*^ cells in the erythroid progenitor population and extensive mis-splicing in clonal hematopoiesis

While *SF3B1* mutations are the most common genetic alterations in MDS patients, they are also associated with a high-risk of malignant transformation in clonal hematopoiesis (CH)^4–8,102,103^. However, the study of *SF3B1* mutations directly in primary human samples has been largely limited to MDS, where confounding co-occurrence of other genetic alterations is common. Thus, CH presents a unique setting to interrogate the molecular consequences of *SF3B1* mutations in non-malignant human hematopoiesis.

We therefore isolated viable CD34+ cells from two CH samples with *SF3B1* mutations (VAFs: 0.15 and 0.22, from CD34+ autologous grafts collected from patients with multiple myeloma in remission) and performed GoT-Splice. A total of 9,007 cells across both samples passed quality filters (**Extended Data Fig. 8a**) and were integrated and clustered based on transcriptome data alone, agnostic to genotyping information (**Fig. 5a; Extended Data Fig. 8b**). Consistent with clinical data indicating normal hematopoietic production, we identified the expected progenitor subtypes using previously annotated progenitor identity markers (**Fig. 5a; Extended Data Fig. 8c, d**). Genotyping data were available for 3,642 cells of these 9,007 cells (40.4%) through GoT (**Extended Data Fig. 9a**). Finally, to exclude additional genetic lesions in these CH samples, we performed copy number analysis with scRNA-seq data and identified no significant chromosomal gains or losses (**Extended Data Fig. 9b**).

**Figure 5.**
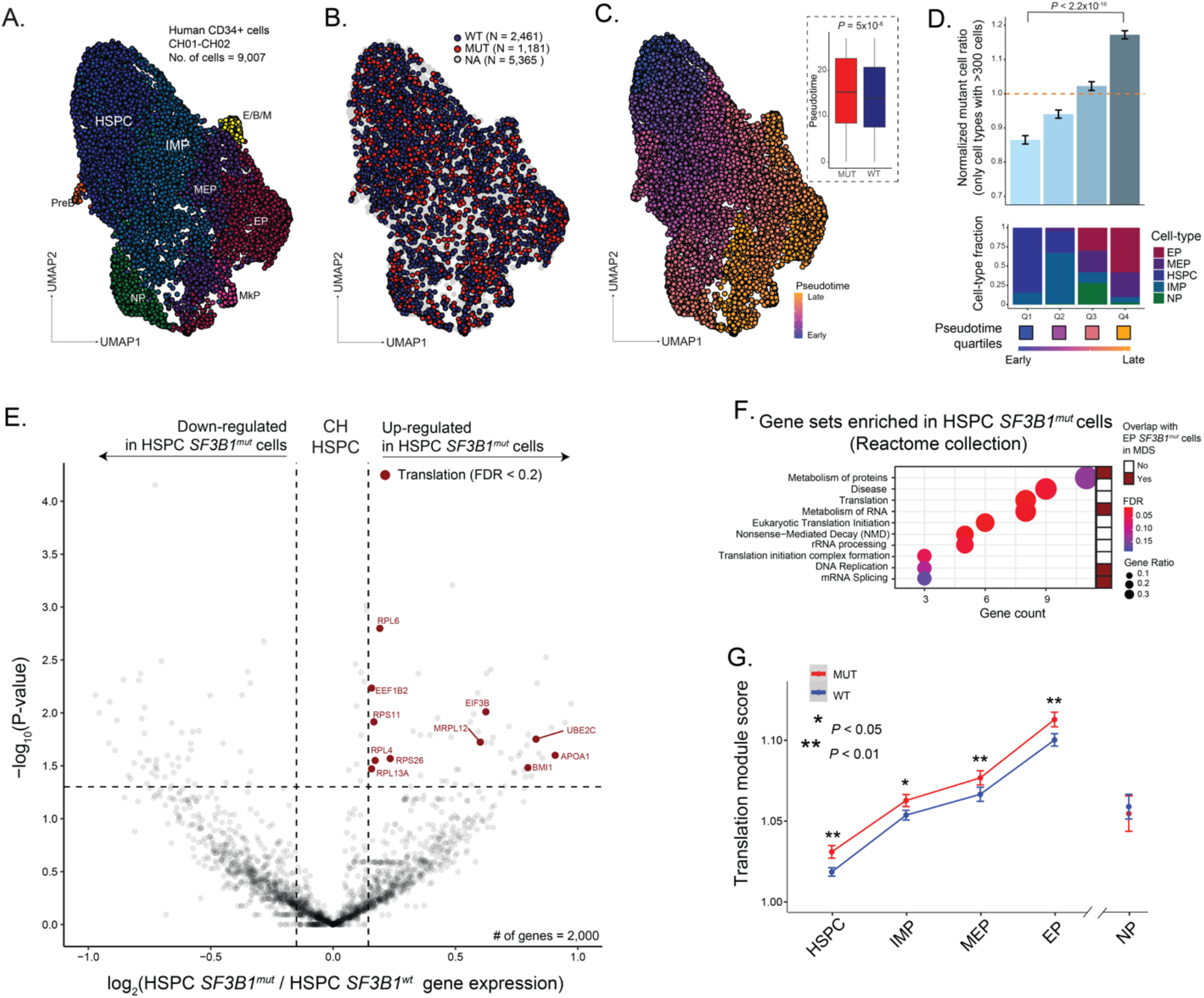
*SF3B1* mutation promotes accumulation of mutant cells along the erythroid lineage in clonal hematopoiesis. **(A)** UMAP of CD34+ cells (*n =* 9,007 cells) from clonal hematopoiesis (CH) samples, one with *SF3B1* K700E mutation and one with *SF3B1* K666N mutation (*n =* 2 individuals), overlaid with cluster cell-type assignments. HSPC, hematopoietic stem progenitor cells; IMP, immature myeloid progenitors; MEP, megakaryocytic-erythroid progenitors; EP, erythroid progenitors; MkP, megakaryocytic progenitors; NP, neutrophil progenitors; E/B/M, eosinophil/basophil/mast progenitor cells; Pre-B, precursors B cells. **(B)** UMAP of CD34+ cells from CH samples overlaid with genotyping data. WT, cells with genotype data without *SF3B1* mutation; MUT, cells with genotype data with *SF3B1* mutation; NA, unassignable cells with no genotype data. **(C)** UMAP of CD34+ cells from CH samples overlaid with pseudotemporal ordering. *Inset:* Pseudotime in *SF3B1*^*mut*^ vs. *SF3B1*^*wt*^ cells in the aggregate of CH01-02. *P*-value for comparison of means from Wilcoxon rank sum test. **(D)** Normalized ratio of mutated cells along pseudotime quartiles. Bars show aggregate analysis of samples CH01-CH02 with mean +/-s.e.m. of 100 downsampling iterations to 1 genotyping UMI per cell. Only cell types with >300 cells were used in the analysis. *P*-value from likelihood ratio test of linear mixed model with or without mutation status. *Bottom*: Fraction of cell types within each pseudotime quartile. **(E)** Differential gene expression between *SF3B1*^*mut*^ and *SF3B1*^*wt*^ HSPC cells in CH samples. Genes with an absolute log_2_(fold change) > 0.1 and *P*-value < 0.05 were defined as differentially expressed (DE). DE genes belonging to the translation pathway (red, Reactome) are highlighted (BH-FDR < 0.2). **(F)** Gene Set Enrichment Analysis of DE genes in *SF3B1*^*mut*^ HSPC cells across CH samples. Gene sets that overlap with *SF3B1*^*mut*^ EP cells in MDS highlighted (red). **(G)** Expression (mean +/-s.e.m.) of mRNA translation-related genes (Reactome) between *SF3B1*^*mut*^ and *SF3B1*^*wt*^ cells in progenitor cells from CH01-02 samples. *P*-values from likelihood ratio test of linear mixed model with or without mutation status.

Projection of the genotyping information onto the differentiation map (**Fig. 5b**), showed no novel cell identities formed by the *SF3B1* mutations, consistent with the fact that patients with CH exhibit no overt peripheral blood count or morphological abnormalities. However, a differentiation pseudotime ordering analysis showed that *SF3B1*^*mut*^ cells are enriched at later pseudotime points when compared to *SF3B1*^*wt*^ cells (**Fig. 5c; Extended Data Fig. 9c**). To further identify differentiation biases in *SF3B1*^*mut*^ CH, we evaluated the mutated cell frequencies across the different prevalent progenitor cell types, as performed in MDS (**Fig. 1e**). Mutated cells were enriched in more differentiated EPs compared to the earlier HSPCs (*P-*value *<* 0.001, linear mixed model, **Fig. 5d; Extended Data Fig. 9d**), showing that *SF3B1*^*mut*^ CH cells already demonstrate an erythroid lineage bias.

To further identify transcriptional dysregulation in *SF3B1*^*mut*^ HSPCs, we performed differential gene expression analysis between mutated and wildtype cells. We observed a similar up-regulation of genes involved in mRNA translation in the *SF3B1*^*mut*^ HSPC in CH (**Fig. 5e, f**; **Supplementary Table 9, 10**), a pathway also observed to be upregulated in our MDS analysis (**Fig. 1h**). In CH, upregulation of mRNA translation pathway genes was observed across multiple cell subtypes along erythroid differentiation, while absent in NPs (**Fig. 5g**). Thus, although no overt blood count abnormalities are observed with *SF3B1* mutation in CH individuals, both the erythroid differentiation bias and aberrant transcriptional profiles are already apparent at this early pre-disease stage.

The analysis of differentially used alternative 3’ splice sites between *SF3B1*^*mut*^ and *SF3B1*^*wt*^ CH cells revealed a marked increase in cryptic 3’ splice site usage in *SF3B1*^*mut*^ cells, as observed in MDS (**Fig. 6a**). These mutant-specific cryptic 3’ splice sites affected genes including *UROD, OXAIL, SERBP1, MED6* and *ERGIC3*, which were also detected to be cryptically spliced in the *SF3B1*^*mut*^ MDS cells. Importantly, the lower VAF associated with pre-malignant CH samples highlights the necessity for GoT-Splice to increase the detection of mis-splicing events occurring at low frequencies, and that may otherwise be missed in bulk sequencing studies (**Fig. 6b; Extended Data Fig. 10a**).

**Figure 6.**
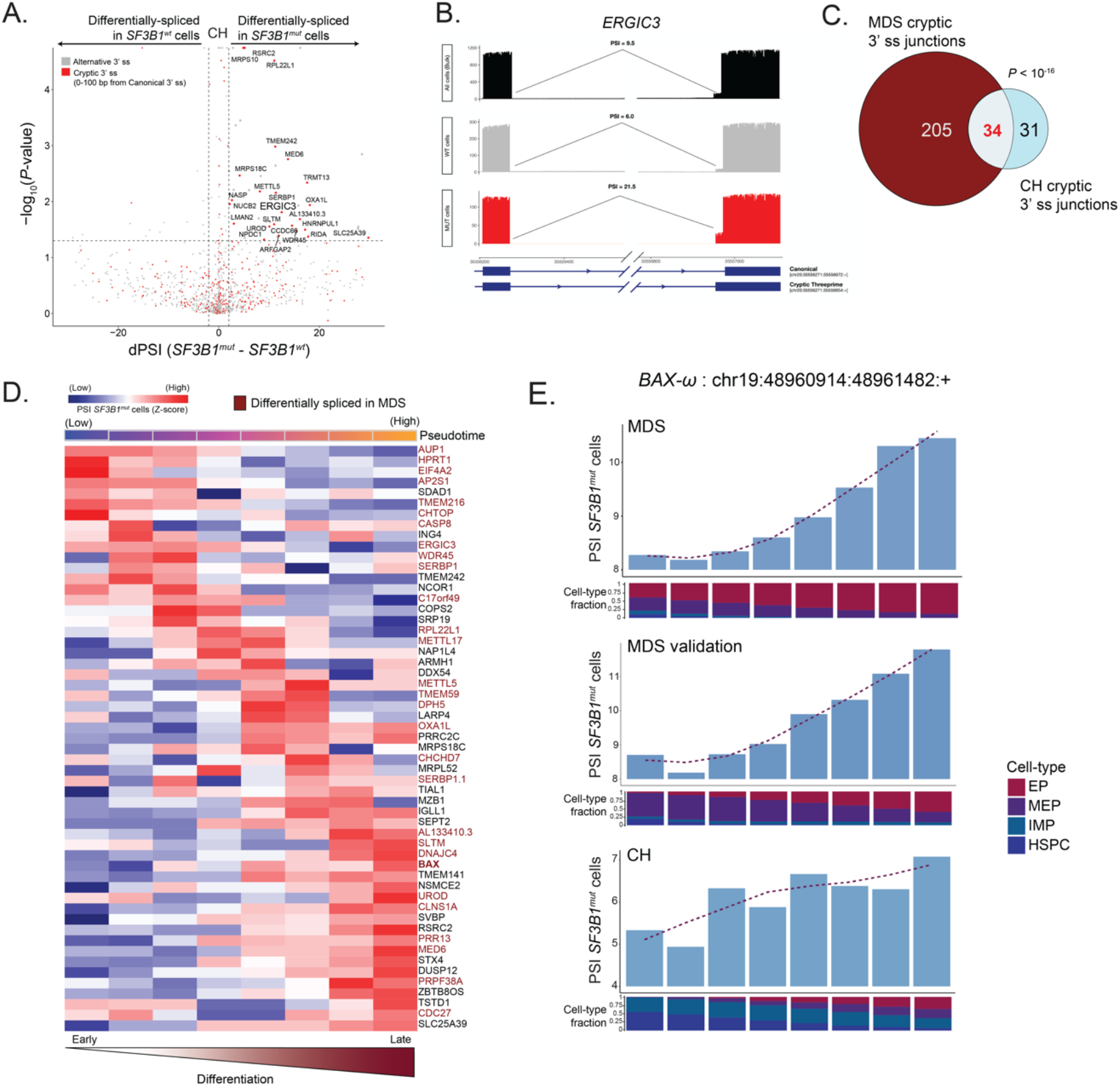
*SF3B1*^*mut*^ clonal hematopoiesis progenitor cells display cell-type specific cryptic 3’ splice site usage. **(A)** Differential splicing analysis between *SF3B1*^*mut*^ and *SF3B1*^*wt*^ cells across CH samples. Junctions with an absolute delta percent spliced-in (dPSI) > 2 and BH-FDR adjusted *P*-value < 0.2 were defined as differentially spliced. **(B)** Sashimi Plot of *ERGIC* intron junction with an *SF3B1*^*mut*^ associated cryptic 3’ splice site showing RNA-seq coverage in *SF3B1*^*mut*^ vs. *SF3B1*^*wt*^ cells within CH samples, as well as compared to the CH samples when treated as bulk (pseudobulk of all cells regardless of genotype). PSI values showing the expected marked increase in the usage of this cryptic 3’ splice site in *SF3B1*^*mut*^ cells alone when compared to both *SF3B1*^*wt*^ cells as well as all cells (pseudobulk of sample). **(C)** Venn Diagram of the overlap of genes with cryptic junctions significantly differentially spliced in at least one erythroid lineage cell type (HSPCs, IMPs, MEPs, EPs) with a dPSI > 2 between MDS01-03 and CH samples. *P*-value for the overlap from Fisher’s Exact test. **(D)** Percent spliced-in (PSI) of junctions in SF3B1^mut^ cells along the hematopoietic differentiation trajectory of erythroid lineage cells. Rows (z-score normalized) correspond to cryptic 3’ splice sites; columns represent the PSI for the usage of a given cryptic 3’ splice site in each window (size of 600 *SF3B1*^*mut*^ cells, sliding by 60 *SF3B1*^*mut*^ cells). Only junctions found to be differentially spliced in at least one cell type with a dPSI > 2 were used in the analysis. Pseudotime across each window shown. Rows are ordered according to the peak in PSI. Cryptic events also found to be differentially spiced in MDS highlighted (red). **(E)** Bar plots of the PSI values for the usage of the *BAX-ω* isoform across each window of *SF3B1*^*mut*^ cells in the MDS, MDS validation and CH cohorts along the hematopoietic differentiation trajectory of erythroid lineage cells. Fraction of cell types in each window shown per cohort (MDS: *SF3B1*^*mut*^ cells (*n =* 6376) ordered by CD71 expression, MDS validation: *SF3B1*^*mut*^ cells (*n =* 987) ordered by pseudotime, CH: MUT cells (*n =* 1021) ordered by pseudotime).

To compare mis-spliced transcripts between CH and MDS, we compared cryptic 3’ splice sites with a *P*-value < 0.05 and dPSI of >= 2 in at least one cell-type along the erythroid differentiation trajectory (HSPC, IMP, MEP or EP) in both CH and in MDS cohorts (**Supplementary Table 11**). While the overall number of significant cryptic 3’ splice sites in CH was lower than in MDS, we observed a significant overlap in shared cryptic events (*P-*value *<* 10^−16^, Fisher’s exact test; **Fig. 6c**). Similarly to MDS, we identified mis-spliced events specific to different stages of erythroid maturation, the majority of which overlapped with MDS cryptic 3’ splice sites (**Fig. 6d**). Notably, CH and MDS showed similar mis-splicing dynamics in the *BAX* transcript along the erythroid differentiation trajectory (**Fig. 6e**).

## DISCUSSION

Here, we present GoT-Splice, a single-cell multi-omics integration that enables joint profiling of genotype, gene expression, protein, and alternative splicing all within the same cell. GoT, as previously described^15^, allows for the comparison between somatically mutated and wildtype cells within the same sample, for genotype to phenotype inferences. Next, by further optimization of long-read sequencing of scRNA-seq libraries^64^, we were able to simultaneously capture both short and long-read data within the same cell, making it possible to analyze the impact of somatic mutations on transcriptional and splicing phenotypes.

To date, few tools are available to process and analyze single-cell long-read data, especially for the purpose of alternative splicing. To address existing analytic gaps, we developed a long-read splicing analysis pipeline that detects and quantifies alternative splicing events within single cells and highlights differential junction usage across cell subpopulations. For processing the long-read data, the pipeline integrates *SiCeLoRe*^64^ to error-correct cell barcodes and UMIs, followed by the generation of consensus reads. Next, unlike other isoform detection methods that perform exon-centric junction calling (such as *SiCeLoRe, TALON*^63^, *FLAME*^104^), we opted for an intron-centric approach followed by split five prime and three prime PSI measurements. Calculating the rate of splicing at the 5’ and 3’ ends of the intron improves the detection of the true splicing rate of each individual intron, compared to exon-centric approaches^68^. In addition, our pipeline detected differential splicing patterns between MUT and WT cells, both across entire samples and within individual cell types, with sample-aware permutation testing to integrate across samples. Finally, the pipeline includes a functional annotation step that provides information regarding the translational consequences of the alternative spliced isoforms. Altogether, our pipeline provides a comprehensive toolkit to process and analyze differential splicing events in scRNA-seq long-read data.

By applying GoT-Splice to the most common splice-altering mutation (*SF3B1)*, we interrogated differentiation biases, differential gene expression, protein expression and splicing patterns, comparing *SF3B1*^*mut*^ vs. *SF3B1*^*wt*^ cells co-existing within the same bone marrow. Importantly, while GoT revealed that *SF3B1*^*mut*^ cells arise early on in uncommitted HSPCs, we observed a differentiation bias of *SF3B1*^*mut*^ cells toward the erythroid progenitor fate. This finding is of particular interest given the clinical association between *SF3B1* mutations and dysplastic erythropoiesis. Differential gene and protein expression in erythroid progenitors revealed signatures that may contribute to this observed differentiation bias of *SF3B1*^*mut*^ cells toward the erythroid fate. Notably, an increase in cell cycle and checkpoint gene expression (*TP53, MDM4* and *CCNE1*) as well as the over-expression of erythroid lineage markers, CD36 and CD71, specifically in *SF3B1*^*mut*^ EPs, suggest a fitness advantage for *SF3B1*^*mut*^ cells along the erythroid lineage.

CH samples likewise showed erythroid biased differentiation with higher mutated cell frequency in committed erythroid progenitors compared with HSPCs. This is one of the first phenotypic studies of clonal mosaicism in human samples, and thus the observation of a somatic mutation-related phenotype, which aligns with the more advanced MDS phenotype, is of particular interest. In our results, *SF3B1*^*mut*^ CH cells showed upregulation of genes in pathways involved in translation and mRNA processing, similar to *SF3B1*^*mut*^ cells in MDS. This finding suggests that the pervasive mis-splicing observed with *SF3B1* mutations may disrupt translation, reminiscent of ribosomopathies, which often also result in dyserythropoiesis^105,106^. Interestingly, it has been shown that overexpression of *MDM4* prevents *TP53* degradation and leads to TP53 complex sequestration, which interferes with p21 activation and results in a sustained cell proliferation. This finding aligns with the observed upregulation of *TP53* and other *TP53*-related pathway genes in *SF3B1*^*mut*^ EPs in MDS. Thus, in addition to the shared erythroid differentiation bias in MDS and CH, aberrant transcriptional profiles linked to a dyserythropoiesis phenotype are also already apparent at the pre-disease CH stage.

Leveraging the single-cell resolution of GoT-Splice and differential splicing analysis between *SF3B1*^*mut*^ and *SF3B1*^*wt*^ cells revealed cell-type specific effects of *SF3B1* mutations on patterns of mis-splicing. First, key genes involved in pathways important for terminal differentiation of hematopoietic stem cells as well as the regulation of erythropoiesis (namely RNA processing, erythroid differentiation, cell cycle and heme metabolism) were found to be cryptically spliced across distinct *SF3B1*^*mut*^ progenitor cell types, many of which were previously reported to be affected in bulk studies of *SF3B1*^*mut*^ MDS^54,72,73^. While some cryptic events were neutral in their effect, many key genes important for erythroid differentiation were found to be NMD-inducing (*e*.*g*., *UROD, GYPA, PPOX*) or cause a frameshift event that may affect protein structure and function (*e*.*g*., BAX) in both the primary and validation MDS cohorts. Thus, our data suggest that mis-splicing of erythroid specific genes and pathways, together with the dysregulation of apoptotic programs, may ultimately lead to the accumulation of *SF3B1*^*mut*^ EPs that fail to reach terminal differentiation^97^, leading to the dyserythropoiesis clinical phenotype. Importantly, this *SF3B1*^*mut*^ mis-splicing phenotype was already evident in the CH samples, suggesting that the impact of somatic CH driver mutations may be conserved from CH to overt myeloid neoplasia.

## Acknowledgments

The work was enabled by the Weill Cornell Flow Cytometry Core. F.G. is supported by an American Society of Hematology Scholar Award (200264-02). N.D. is supported by a F30 Predoctoral Fellowship from the NHLBI of the National Institutes of Health (F30HL156496) and by a Medical Scientist Training Program grant from the National Institute of General Medical Sciences of the National Institutes of Health under award number T32GM007739 to the Weill Cornell/Rockefeller/Sloan Kettering Tri-Institutional MD-PhD Program. R.C. was supported by Lymphoma Research Foundation and Marie Skłodowska-Curie fellowships. D.A.K. is supported by NSF CAREER award DBI2146398. D.H.W. is supported by the Oglesby Charitable Trust. D.A.L. is supported by the Burroughs Wellcome Fund Career Award for Medical Scientists, Valle Scholar Award, the National Institutes of Health Director’s New Innovator Award (DP2-CA239065), Leukemia Lymphoma Scholar Award and the Mark Foundation Emerging Leader Award. This work was supported by the National Heart Lung and Blood Institute (R01HL157387-01A1 and R01HL128239), the National Cancer Institute (R01s CA242020, R01 CA251138, and P50 254838), the Edward P. Evans MDS Foundation, a Tri-Institutional Stem Cell Initiative award and the National Human Genome Research Institute, Center of Excellence in Genomic Science (RM1HG011014). This work received research support from Oxford Nanopore Technologies.

## Author contributions

F.G., P.C., A.G.H., M.C.-L., J.T., O.A.-W., D.A.L. conceived the project and designed the study. K.B., D.H.W, I.M.G., O.A.-W. provided clinical samples. A.G.H., A.D.S., S.G., X.D., S.H., R.C. performed the experiments. F.G., P.C., A.G.H., M.C.-L., L.K., C.C., J.B., A.W.D., N.D., G.M., J.S. performed the analyses. F.G., P.C., A.G.H., M.C.-L., T.H.A., R.C., S.J., E.H., D.A.K., I.M.G., J.T., O.A.-W, D.A.L helped interpret results. F.G., P.C., A.G.H., M.C.-L., O.A.-W., D.A.L. wrote the manuscript. All authors reviewed and approved the final manuscript.

## Code availability

IronThrone v.2.1 pipeline is available at https://github.com/landau-lab/IronThrone-GoT. ONT Single-Cell Splice pipeline is available at https://github.com/landau-lab/ONT-sc-splice.

## Data Availability

The raw FASTQ files are being deposited to the European Genome-Phenome Archive.

## METHODS

### Patient samples

The study was approved by the local ethics committee and by the Institutional Review Board (IRB) of Weill Cornell Medicine, University of Manchester and Dana Farber Cancer Institute, conducted in accordance with the Declaration of Helsinki protocol. Cryopreserved mononuclear cells isolated from bone marrow biopsies or peripheral blood from myelodysplastic syndrome patients with *SF3B1* mutations were retrieved from Memorial Sloan Kettering and University of Manchester. Additionally, cryopreserved G-CSF mobilized stem cell grafts (without additional mobilizing agents such as plerixafor or cyclophosphamide) from CH patients with *SF3B1* mutations were retrieved from the Dana Farber Cancer Institute (**Supplementary Table 1**). Cryopreserved mononuclear cells and grafts were thawed and stained using standard procedures. Cells were first incubated with Human FcX blocking solution (Biolegend, #422302) and then incubated with the surface antibody CD34-PE-Vio770 (clone AC136, lot #5180718070, dilution 1:50, Miltenyi Biotec) and DAPI (Sigma-Aldrich) for 10 minutes at 4°C. Cells were then sorted for DAPI-negative, CD34+ cells using BD Influx at the Weill Cornell Medicine flow cytometry core.

### GoT-Splice with CITE-seq

GoT-Splice with CITE-seq integrates Genotyping of Transcriptomes (GoT) with both long-read single-cell transcriptome profiling (with Oxford Nanopore Technologies [ONT]) and proteogenomics (with CITE-seq). GoT was performed as previously described^15^. For samples without CITE-seq, CD34+ cells were sorted, and RNA was prepared for sequencing following the standard 10x Genomics Chromium 3’ (v.3.1 chemistry) protocol and according to manufacturer’s recommendations for the generation of scRNA-seq libraries (**Fig. 1a**). For GoT-Splice samples that were processed with CITE-seq, prior to sorting, cells were blocked with FcX block for 15 minutes prior to being stained with Total-SeqA antibodies for 30 minutes on ice (see **Supplementary Table 3** for list of antibodies used). The standard 10x Genomics Chromium 3’ (v.3.1 chemistry) and CITE-seq protocols^35,36^ were carried out according to manufacturer’s recommendations for the generation of scRNA-seq and ADT libraries (**Fig. 1a**). At the cDNA amplification step in the 10x Genomics protocol, 1 μL of 1 μM spike-in primer (5’-GATCCTCGTCCTCATTGAACCGC-3’) was added to increase the yield of *SF3B1* cDNA and 1 μL of 0.2 μM ADT PCR additive primer (5’ – CCTTGGCACCCGAGAATTCC – 3’) was added to amplify ADT. After cDNA amplification and a double-sided cleanup with SPRI beads to separate cDNA and ADT fractions, the ADT fraction was amplified for 10 cycles with SI-PCR oligo (10x Genomics) and TruSeq Small RNA RPI-x (Illumina) primers to index the samples. SPRI was used to clean up the ADT final products. In both samples in which CITE-seq was conducted and not conducted, cDNA was allocated for gene expression library creation (standard 10x protocol; 25% of cDNA), targeted genotyping (10% of cDNA), and ONT sequencing with biotin enrichment (10 ng of cDNA). Any remaining cDNA was stored. For locus-specific amplification (GoT), two serial PCRs were performed with nested reverse primers, based on the *SF3B1* mutation of interest. For mutations upstream of K700E, (5’-GATCCTCGTGGTCATTGAACCGC-3’ and 5’-CACCCGAGAATTCCAGGCTACTATGATCTCTACCATGA GACCTG-3’) and, for K700E mutations, (5’-GTGCAAAAGCAAGAAGTCCT-3’ and 5’-CACCCGAGAATTCCATGAACATGGTCTTGTGGATGAG-3’) were used as reverse primers. These reverse primers and the generic forward SI-PCR amplify the site of interest from the cDNA template (10 PCR cycles each). The second locus-specific reverse primers contain a partial Illumina TruSeq Small RNA read 2 handle and a locus-specific region to allow *SF3B1* specific priming. The SI-PCR oligo (10x Genomics) anneals to the partial Illumina TruSeq read 1 sequence, preserving the cell barcode (CB) and unique molecule identifier (UMI). After these rounds of amplification and SPRI purification to remove unincorporated primers, a third PCR was performed with a generic forward PCR primer (P5_generic, 5’ – AATGATACGGCGACCACCGAGATCTACAC – 3’) to retain the CB and UMI together with an RPI-x primer (Illumina) to complete the P7 end of the library and add a sample index (6 PCR cycles).Gene expression, ADT, and *SF3B1* amplicon libraries were pooled to receive 25,000, 5,000, and 5,000 reads per cell, respectively, during Illumina sequencing. The cycle settings were as follows: 28 cycles for read 1, 90 cycles for read 2, 10 cycles for i7, and 10 cycles for i5 sample index. To examine splicing patterns broadly in the whole transcriptome, full length cDNA was sequenced using the Oxford Nanopore Technologies sequencing on PromethION and GridION flow cells. To enrich for transcripts that contain CBs and UMIs and decrease the presence of PCR artifacts, on-bead PCR with a biotinylated primer selecting for an adapter upstream of the CB was completed^64^ (**Fig. 2a**). In brief, 10 ng of full-length cDNA was amplified with LongAmp master mix (NEB) and TSO (5’-NNNAAGCAGTGGTATCAACGCAGAG-3’) and biotinylated read 1 (5’-/5Biosg/AAAAACTACACGACGCTCTTCCGATCT-3’) primers for 5 cycles. M270 streptavidin beads (ThermoFisher) were washed with 1X SSPE buffer, resuspended in 5X SSPE buffer and incubated with PCR amplicon after clean up with 0.8X SPRI beads. After a 15-minute incubation, the beads were washed with 1X SSPE and 10 mM Tris-HCl (pH 8) resuspended in PCR master mix, and further amplified with LongAmp master mix, TSO and read 1 (5’ – NNNCTACACGACGCTCTTCCGATCT – 3’) primers for 5 cycles. After cleanup with SPRI, 100-300 ng of each full-length cDNA library was sequenced on one PromethION or GridION flow cell with SQK-LSK110.

### ScRNA-seq Illumina data processing, alignment, clustering and cell-type classification

10x Illumina data was processed using Cell Ranger (v.3.1.0) with default parameters and reads were aligned to the human reference sequence GRCh38. For all samples, the Seurat package (v.3.1) was used to perform QC filtering, and unbiased clustering of CD34+ sorted cells^107^. As an overview, for each sample dataset, cells with number of UMIs (nCount_RNA) <1000 or nCount_RNA > 3 s.d. above the mean nCount_RNA value, number of unique genes (nFeature_RNA) > 3 s.d. above the mean nFeature_RNA value and mitochondrial gene percentage (perc.mito) > 20% were filtered. Using the SCTransform function, each dataset was log normalized using the default scale factor of 10,000, scaled and potential confounders (such as nCount_RNA, perc.mito and S phase and G2M phase gene expression scores) were regressed out of the data. SCTransform also identified the top 3000 variable genes found in each dataset that are used for integration. Before clustering, the individual datasets were integrated based on disease status (i.e. primary MDS samples, MDS01-03, were integrated together, MDS validation samples, MDS04-06, from patient treated with growth factors at the time of biopsy were integrated together and then the CH samples, CH01-02, were integrated together) and underwent batch correction within Seurat which implements canonical correlation analysis (CCA) and the principles of mutual nearest neighbors (MMN)^108^. For integration, 30 canonical vectors were used for the CCA in the FindIntegrationAnchors function, and 30 principal components were used for the anchor weightings step in the IntegrateData function (as recommended in Seurat). Next, a principal component analysis (PCA) was performed using the variable genes of the integrated dataset and the JackStraw method was used to determine statistically significant principal components (PCs) to be used as inputs into the UMAP algorithm for cluster visualization. Clustering was performed with the FindNeighbors (using only significant PCs) and the FindClusters (resolution = 2) functions which rely on the k-nearest neighbors (KNN) algorithm to identify cell clusters. Unique clusters were manually assigned on the basis of differentially expressed genes identified with the FindAllMarkers function which looked only at genes found in at least 25% of cells in either of the two input comparison groups and only returned results for genes with at least a 0.25 log transformed fold change between groups. More specifically, cluster annotations were made according to the differential expression of canonical lineage marker genes identified in previous single-cell RNA-seq data of normal hematopoietic progenitor cells^37^ (**Supplementary Table 2**). Clusters with similar increased expression of these canonical markers were merged to form the main progenitor subsets: HSPCs, IMPs, NPs, MkPs, MEP, EPs, Pre-Bs and E/B/Ms in the primary MDS, MDS validation and CH cohort as well as Mono, MonoDCs, DCs, B cells and T cells in MDS and MDS validation. Finally, pseudotime analysis was performed using the Monocle3 R package with recommended parameters (v.0.2.1)^109^.

### IronThrone GoT for processing targeted amplicon sequences and performing mutation calling

Genotyping of single cells was carried out with the IronThrone (v.2.1) pipeline as previously described^15,110^. In brief, individual amplicon reads were assessed for the appropriate structure (*i*.*e*., presence of the primer sequence and the expected sequence between the primer and given mutation site) and all reads were assessed for a matching cell barcode to the list generated from the 10x paired GEX dataset. A Levenshtein distance of 0.1 was allowed for all sequence matching and collapsing steps and only UMIs with a minimum of 2 supporting reads were retained for final genotyping. Following UMI collapse, genotype assignment of individual UMIs was conducted as described previously with majority rule of supporting reads for wildtype or mutant status (using a 0.7 PCR read ratio, above which the majority of PCR reads must be in order for a UMI to be called definitively). Rare UMIs that did not pass this threshold were removed as ambiguous. Additionally, to remove reads that result from PCR recombination, UMIs in the amplicon library that match UMIs of non-*SF3B1* genes in the gene expression library were discarded. Finally, given the heterozygous nature of these *SF3B1* mutations, each single cell was assigned as either mutant (MUT) or wildtype (WT) as follows: cells with at least 1 mutant UMI were assigned as MUT cells and cells with 0 mutant UMIs and at least 1 wildtype UMI were assigned as WT. While the genotyping information is derived from transcribed molecules alone and may be affected by whether transcripts from wildtype versus mutant alleles were expressed and/or captured, the fraction of MUT cells as determined by GoT using all cells with at least 1 UMI yielded similar values to those determined by bulk DNA exon sequencing (**Extended Data Fig. 4a**). Despite this, we systematically applied specific approaches to exclude the effect of this confounder (that is, the expression level of the target gene) on the conclusions of other downstream analyses. Firstly, to rule out the possibility that higher *SF3B1* expression results in a greater ability to detect mutant alleles, and thereby in a higher mutant-cell frequency, we downsampled all cells to a single amplicon UMI before mutation calling when conducting the mutant-cell frequency analyses. Finally, for the remaining of our downstream analyses between *SF3B1* mutant and wildtype cells (except for the differential gene expression and gene set enrichment analyses in CH due to low numbers of genotyped cells), we took the more conservative approach considering only genotyped cells with two or more genotyping amplicon UMIs. As benchmarking, the *SF3B1* genomic regions of interest that were used for GoT were examined in each matching GEX library to determine how many UMIs were able to successfully capture the targeted sequence in conventional 10x data (**Extended Data Fig. 4b, 9a**).

### Mutant cell frequency

The frequency of mutant cells, as determined by GoT, was assessed as previously performed in Nam et al.^110^. Firstly, we used only cells with at least 1 UMI and only considered cell types with at least 300 genotyped cells. To account for the potential confounding effect of a heterozygous mutation as well as variable *SF3B1* expression, we performed amplicon UMI down-sampling to 1 UMI per genotyped cell prior to mutation calling for calculating MUT cell frequencies. An equal number of cells from each sample within the MDS cohort, were subsampled randomly for the integrated data to ensure equal representation from each patient. Genotyping amplicon UMIs were downsampled (x100 iterations) to 1 UMI per cell and MUT cell frequency was determined for each progenitor cluster for either the integrated dataset or individual samples. This frequency was then divided by the total mutant cell frequency across all progenitor subsets for each of the iterations. Linear mixed effects analysis was performed using the lme4 package (v.1.2-1). Progenitor identity was defined as the fixed effect, and for random effects, we used intercepts for individual patients (subjects) and iterative downsampling. *P*-values were obtained by likelihood ratio tests of the full model with the fixed effect against the model without the fixed effect^111^.

### Differential gene expression and gene set enrichment

The differential gene expression analysis (DGEA) comparing WT and MUT cells and gene set enrichment analysis (GSEA) were performed as done in Nam et al.^110^. In brief, for each cohort we used a within-sample permutation test for the analysis of each progenitor cell subtype. To ensure equal representation from each patient, we downsampled the total number of mutated and wildtype cells to the same number across all patients. The observed log_2_ fold change values were calculated comparing the MUT versus WT cells for the tested genes. The tested genes included the top 2,000 most variable genes (excluding mitochondrial genes) which were filtered for those expressed in at least 10% of either group (MUT versus WT), for each progenitor subtype. Next, the WT and MUT labels were shuffled over 100,000 iterations, within each patient, and fold change values were re-calculated to create a background distribution. *P*-values were calculated per gene as a percent of permutations whose absolute fold change values were more extreme than the absolute value of the observed fold change (**Supplementary Table 5, 9**). Hypergeometric test for GSEA of the integrated differentially expressed genes (*P*-value < 0.05, log_2_(fold change) > 0.1) was performed using the Cluster Profile package (v. 0.1.9). FDR multiple hypothesis testing correction was performed. MSigDB C2 curated gene sets were included in the analyses (**Supplementary Table 6, 10**).

### ADT processing

CITE-seq was performed on the primary MDS cohort (for samples MDS02-03) and as mentioned above, the 10x Illumina ADT data was processed using Cell Ranger (v.3.1.0) with default parameters and counts were generated for each marker in the CITE-seq panel (**Supplementary Table 3**). After using the Seurat package (v.3.1) for QC filtering, and unbiased clustering of the CD34+ sorted cells based on RNA data, ADT data was also normalized using centered log-ratio (CLR) normalization, scaled and the expression of various ADT markers was used in confirming the cell-type assignment of different progenitor subsets. For benchmarking purposes, Seurat’s Weighted Nearest Neighbor (WNN) Analysis was also performed, which is a multi-modal analysis that integrates both RNA and ADT data when performing cell clustering. This was used to compare to the clustering output when using the KNN algorithm that relies on RNA data alone (**Extended Data Fig. 2d**). For the WNN analysis, cells were filtered and integrated using SCTransform (as described above). The RNA data was logNormalized and the ADT data was run through CLR normalization and the RunPCA function for dimensionality reduction was also run independently on each modality. Next, the FindMultiModalNeighbors function was used which for each cell, calculates its closest neighbors in the dataset based on a weighted combination of RNA and ADT similarities. This constructs a WNN graph that was visualized with the RunUMAP function. The cell-type assignments generated from the initial clustering (with RNA data alone) were then projected onto this new UMAP for comparison (**Extended Data Fig. 2d**).

### Denoised scaled by background normalization (DSB) filtering and differential protein expression

We used the dsb package^112^ (v.0.1.0) as an alternative form of normalization for the ADT protein expression values. Normalized values were applied for selection filtering of ADT markers for which the true signal was above the background noise levels, within the captured cell-contained droplets. dsb discriminates between background noise by differentiating between empty droplets (containing ambient mRNA and antibody but no cell) and true cell-containing droplets. The background matrix was defined from the comparison of the raw feature barcode matrices from the 10x sequencing output versus the processed filtered feature barcode matrix results generated from running Cell Ranger (see Methods above). The final output filters out empty droplets and retains only true cell-containing droplets based on the 10x cell calling algorithm. As such, the matrix of background noise is generated by subtracting out the positive cell containing droplets found in the filtered matrices from the negative empty droplets in the raw matrices. Furthermore, with an additional filter requiring the removal of drops with protein library size > 1.5 and number of genes < 80 was applied to refine the background noise signal. Normalization was performed using the DSBNormalizeProtein function omitting isotype controls and denoised counts. The dsb normalized values were defined as the number of standard deviations above the background noise and antibodies were then filtered, keeping only those with a dsb normalized expression value of > 2 in at least 1 cell-type (**Supplementary Table 4**).

When performing the differential protein expression analysis across our patient samples, we used an iterative downsampling (x1,000) approach that, at each iteration, randomly samples an equal number of *SF3B1*^mut^ and *SF3B1*^wt^ cells from each patient sample before calculating the median log_10_FC of protein expression between *SF3B1*^mut^ / *SF3B1*^wt^ cells. This was done to ensure equal representation of genotyped cells from each patient. To calculate the median log_10_FC of *SF3B1*^mut^ / *SF3B1*^wt^ cells, we first modified the Seurat’s FindMarker function to calculate the median instead of the mean expression, a measure that is more robust to outlier values. Then, for each downsampled object we obtained a table containing the log_10_FC of each antibody per cell-type. log_10_FC matrices are combined by taking the median across the downsampled iterations, resulting in the median log_10_FC values. Statistical significance was assessed by performing permutation tests (x10,000) within each patient sample matrix. (**Fig. 1f**) shows normalized ADT expression across cell-types, using the maximum expression.

### ScRNA-seq ONT long-read sequencing data processing, alignment, junction calling and annotation

Guppy (v.3.0.6 -4.0.11) was used for base calling FAST5 files output from ONT sequencing. We then filtered for only reads containing a polyA tail within 100 base pairs of either 5’ or 3’ end using the ‘NanoporeReadScanner-0.5.jar’ within the SiCeLoRe-1.0 workflow. Filtered reads are aligned to the primary human genome, assembly GRCh38.p12 using minimap2 (v.2.17). Minimap2 was used with the ‘-ax splice’ flag to prioritize annotated splice junctions. Additionally, we made use of the ‘--junc-bed’ option, to increase alignment scores for those splice junctions found in the reference junction bed file. For our reference junctions, we used splice junctions from single-cell SMART-seq2 data from human CD34+ cells obtained from a CH sample with no *SF3B1* mutation. Additionally, we used ‘--secondary=no’ to suppress multi-mappings. In preparation to identify the cell barcodes and UMIs present in the long-read sequencing, we used the ‘IlluminaParser-1.0.jar’ in SiCeLoRe to parse the cell barcodes and UMIs present in the complementary short-read sequencing library. We continued to use SiCeLoRe to tag the aligned BAM files with cell barcodes and UMIs identified in the short-read library and generate consensus sequences for each unique cell barcode and UMI combination. Consensus sequences were used to create a gene by cell count matrix. For all other steps, we used the default parameters set by SiCeLoRe-1.0, following the workflow found at https://github.com/ucagenomix/sicelore. Intron-junction calling is then performed on consensus sequence BAM files, adapted from the method used in the LeafCutter pipeline for short-read RNA-seq data^69,113^. In brief, the intron-junction calling pipeline utilizes the pysam.fetch() function and iterates through each transcript in the BAM file, noting its cell barcode (CB) tag as well as the coordinates of each intron-junction for that transcript. On iterating through the BAM file, counts for the usage of each unique intron-junction and the corresponding CB are recorded. This ultimately generates an Intron-Junction x Cell Barcode count matrix for the given BAM file. Each intron-junction is then identified using annotations available in the GENCODE GRCh38.p12 v31 basic annotation reference file as either canonical 3’, canonical 5’, alternative 3’, alternative 5’. This outputs a metadata file with annotations for each junction corresponding to the junctions of the Intron-Junction x Cell Barcode count matrix. The metadata included the 3’ and 5’ sites defining each junction, the distance from the canonical 3’ or 5’ site end for each start and end site, and the classification of each site. Alternative 3’ and 5’ junctions were further broken down into alternative and cryptic based on the distance the junction was from the canonical splice site. If the alternative splice site was within 100 base pairs of the canonical splice site, it was classified as a cryptic splice site. Given the intron-centric approach of the pipeline, each event could be classified as either annotated, alternative 3’, alternative 5’, a cryptic 3’ splice site, a cryptic 5’ splice site or an exon-skipping event (see Methods below for exon-skipping annotation; **Supplementary Table 7, 8, 11**).

### Differential transcript usage

All alternative 3’ junctions were filtered to only include those that contained at least 5 total reads. To identify differentially used transcripts between *SF3B1*^*mut*^ and *SF3B1*^*wt*^ cells, junction reads were then pseudobulked based on mutation status across all MDS patients or all CH patients. We then computed the log_10_(odds ratio) of the likelihood of each junction being observed in the MUT cells over the WT cells. The genotype labels of each of the cells was permuted 100,000 times and we then repeated pseudobulking and computation of the log(odds ratio) of each junction. Permutations of the genotype label were patient aware, so the mutant cell frequency across patients was unchanged for each permutation. The *P*-value was determined based on the likelihood of seeing the observed odds ratio in comparison to the null distribution of the permuted odds ratios for each junction. The same testing was done within each cell-type to identify the differentially used junctions between *SF3B1*^mut^ and *SF3B1*^wt^ cells within a specific cell-type. We classified junctions as differentially spliced events if they had *P-*value *<* 0.05 and delta percent spliced in (dPSI) of >= 2 (a positive dPSI here represents a splicing event more highly used in the *SF3B1*^*mut*^ population of cells). To observe the usage of these differentially spliced cryptic 3’ events (*P-*value *<* 0.05 and dPSI >= 2) across the continuum of erythroid maturation as opposed to within discrete cellular states, erythroid lineage MUT cells (HSPCs, IMPs, MEPs and EPs) were ordered from least to most differentiated, grouped into bins and the MUT cell PSI for each cryptic event was calculated per bin (**Fig. 4a, 6d**). Specifically, in the primary MDS cohort (**Fig. 4a**), 6301 *SF3B1*^*mut*^ cells were ordered by the expression of the erythroid marker CD71 (obtained from CITE-seq) and a bin size of 3000 *SF3B1*^*mut*^ cells, sliding by 300 *SF3B1*^*mut*^ cells at each step, was used to capture the continuous change in the usage of the different cryptic 3’ junctions via the MUT cell PSI measurements per bin. The variance in the usage of each cryptic 3’ event was measured by calculating the range of PSIs across all the bins along the continuum and only cryptic junctions that had a PSI range of at least 2 and average coverage across all bins of 10 reads were considered. This approach was taken to focus on cryptic events that had a variable signal and that were also well supported. In CH (**Fig. 6d**), 1020 MUT cells were ordered by pseudotime and a bin size of 600 *SF3B1*^*mu*t^ cells, sliding by 60 *SF3B1*^*mut*^ cells at each step, was used to capture the MUT cell PSI per bin. Similarly, only cryptic junctions that had a PSI range of at least 2 and average coverage of 10 reads were considered.

For the *BAX* cryptic event (**Fig. 6e**), in order to directly compare the per bin PSI values across all 3 cohorts (MDS, MDS validation and CH) we adjusted the bin and window sizes across the cohorts to ensure the same number of final bins for each cohort. In order to achieve this, we took the following approach: MDS - MUT cells ordered by CD71 expression, window size of 3750 *SF3B1*^mut^ cells, sliding by 375 *SF3B1*^*mut*^ cells, MDS validation: cells ordered by pseudotime, window size of 580 *SF3B1*^*mut*^ cells, sliding by 58 *SF3B1*^*mut*^ cells, CH: cells ordered by pseudotime, window size of 600 SF3B1^mut^ cells, sliding by 60 *SF3B1*^*mut*^ cells. To note, for each of the sliding window analyses, only MUT cells with at least 2 genotyping amplicon UMIs were considered.

### Exon skipping and nonsense-mediated decay (NMD) annotations

In order to identify exon skipping events, for each gene in the GENCODE GRCh38.p12 v31 basic annotation reference file we determined its main functional isoform (as those that belong to the APPRIS database and carry the “appris_princial” tags) to compare to the transcript isoforms generated in our data. With this, for a given gene, each identified intron junction within our data was compared to the reference and labeled as an “exon_skip” if it excluded any of the exons present in the reference. The number of exons skipped was also recorded. To identify NMD inducing alternative splicing events, we also developed a pipeline that inspects each intron junction in the Intron-Junction x Cell Barcode count matrix and detects the presence of premature termination codons (PTCs) and frameshift events induced as a result of alternative splicing. In brief, this is done by grabbing the entire nucleotide sequence of a particular isoform noting the position of the last exon-exon junction, finding the position of the first start codon and from there, phasing along the triplets of nucleotides of that given sequence string. By following the known rules of NMD, each intron junction was further annotated as being NMD-inducing (which would lead to NMD of its associated transcript) or NMD-neutral. Specifically, the 50-nucleotide rule was followed such that an event is labeled NMD-inducing if a PTC is introduced greater than 50 nucleotides away from the last exon-exon junction or NMD-neutral if a PTC is introduced within 50 nucleotides of the transcript’s last exon-exon junction. Finally, Intron-junctions were labeled to cause frameshifts if the total number of nucleotides involved in an alternative 3’ or 5’ splicing event was not divisible by 3. Altogether, each intron junction was classified as follows: (i) NMD-inducing event (due to the introduction of a PTC); (ii) NMD-neutral with a frameshift event; and (iii) NMD-neutral with no frameshift event.

### Motif enrichment analysis

High quality cryptic 3’ junctions (MUT read coverage > 3, PSI >= 2, junction cluster read coverage > 20 across at least 2 junction clusters) were obtained from the junction quantification matrix from samples MDS05-06. Each of these cryptic 3’ splice sites were then paired to a corresponding canonical junction, requiring both, canonical and cryptic junctions, to be part of the same splicing cluster (as described above). Flanking sequences, 50 nucleotides upstream and 10 nucleotides downstream of the 3’ splice site were obtained from the two junction sets and used to calculate position weight matrices (PWM). For each position, a log odds ratio enrichment for each nucleotide was calculated using Fisher’s exact test, comparing the cryptic 3’ splice site nucleotide composition against the canonical. Reported positions were filtered according to their enrichment significance (*P-*value *<* 0.05).

## EXTENDED DATA

**Extended Data Figure 1.**
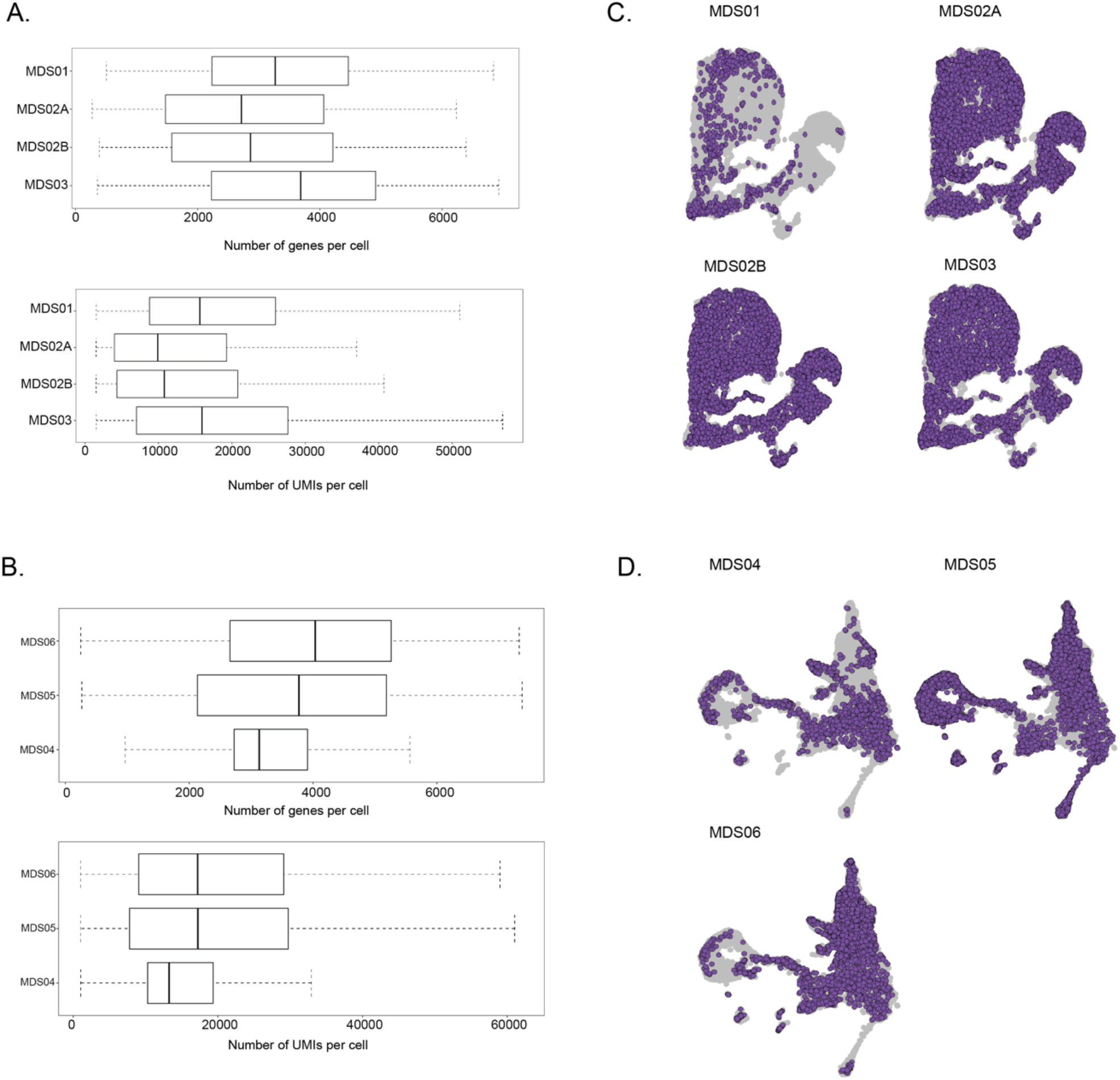
MDS and MDS validation QC and integration. **(A)** Number of genes per cell (top) and number of UMIs per cell (bottom) in CD34+ sorted hematopoietic progenitors from samples MDS01-03 after QC filters, shown by each patient sample. **(B)** Number of genes per cell (top) and number of UMIs per cell (bottom) in CD34+ sorted hematopoietic progenitors from samples MDS04-06 after QC filters, shown by each patient sample. **(C)** UMAP of CD34+ sorted progenitor cells for each individual sample of MDS01-03 after integration using the Seurat package. **(D)** UMAP of CD34+ sorted progenitor cells for each individual sample of MDS04-06 after integration using the Seurat package.

**Extended Data Figure 2.**
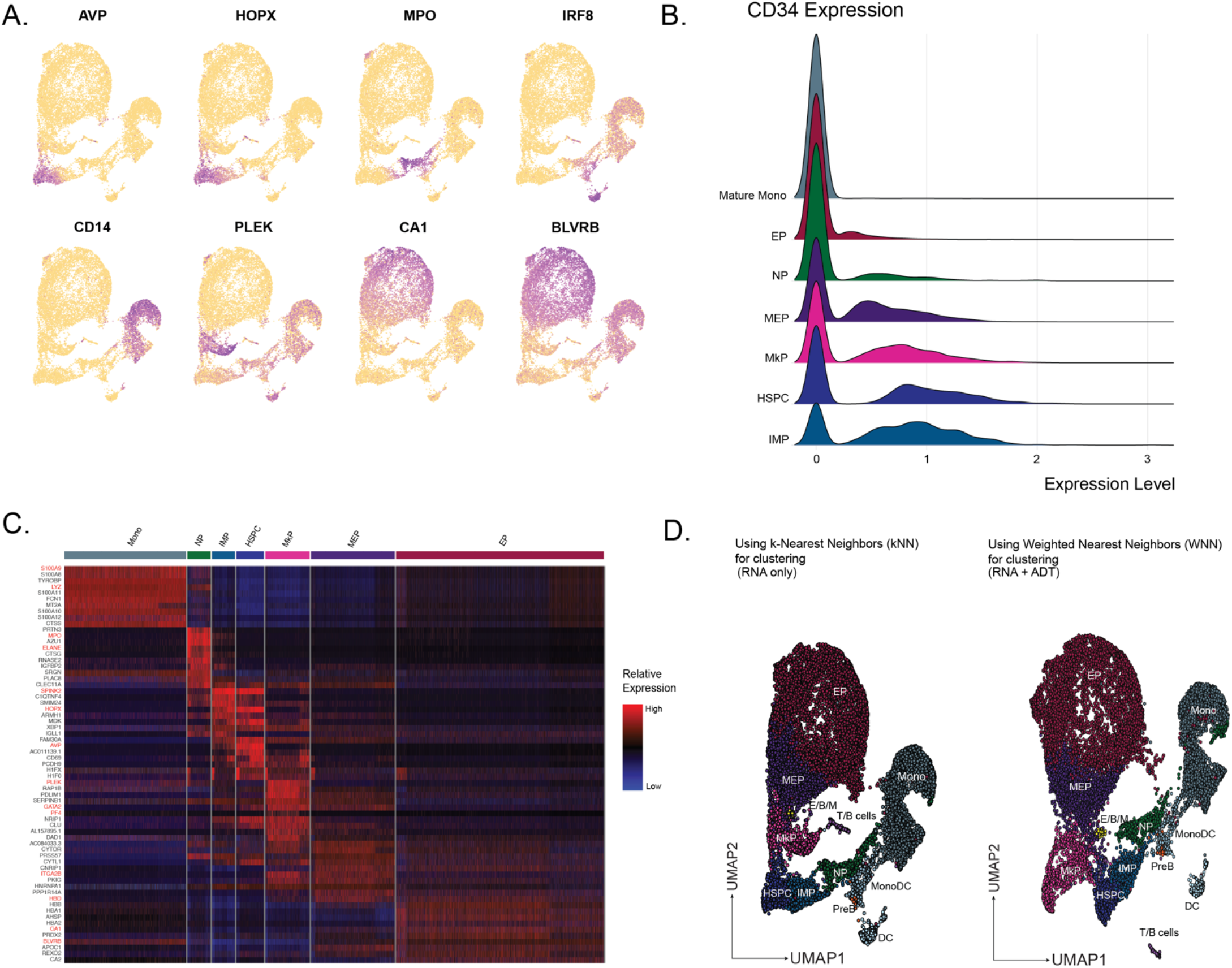
MDS clustering and cell-type assignment. **(A)** Expression of lineage-specific genes from Velten et al.^37^ scored and projected onto the UMAP representation of cells from MDS01-03. **(B)** CD34 expression per progenitor cell-type of CD34-monocytes among CD34+ sorted hematopoietic progenitors. **(C)** Heatmap of top 10 differentially expressed genes for each progenitor subset for MDS01-03. **(D)** UMAPs comparing the graph-based clustering output when using the k-nearest neighbors (KNN) algorithm to perform clustering of cells with RNA data alone vs. when using the weighted nearest neighbors (WNN) algorithm that allows for the integration of both RNA and ADT data for clustering cells. The cell-type assignments determined from the KNN-RNA based clustering are projected onto the WNN-RNA+ADT clustering for comparison between the two methods.

**Extended Data Figure 3.**
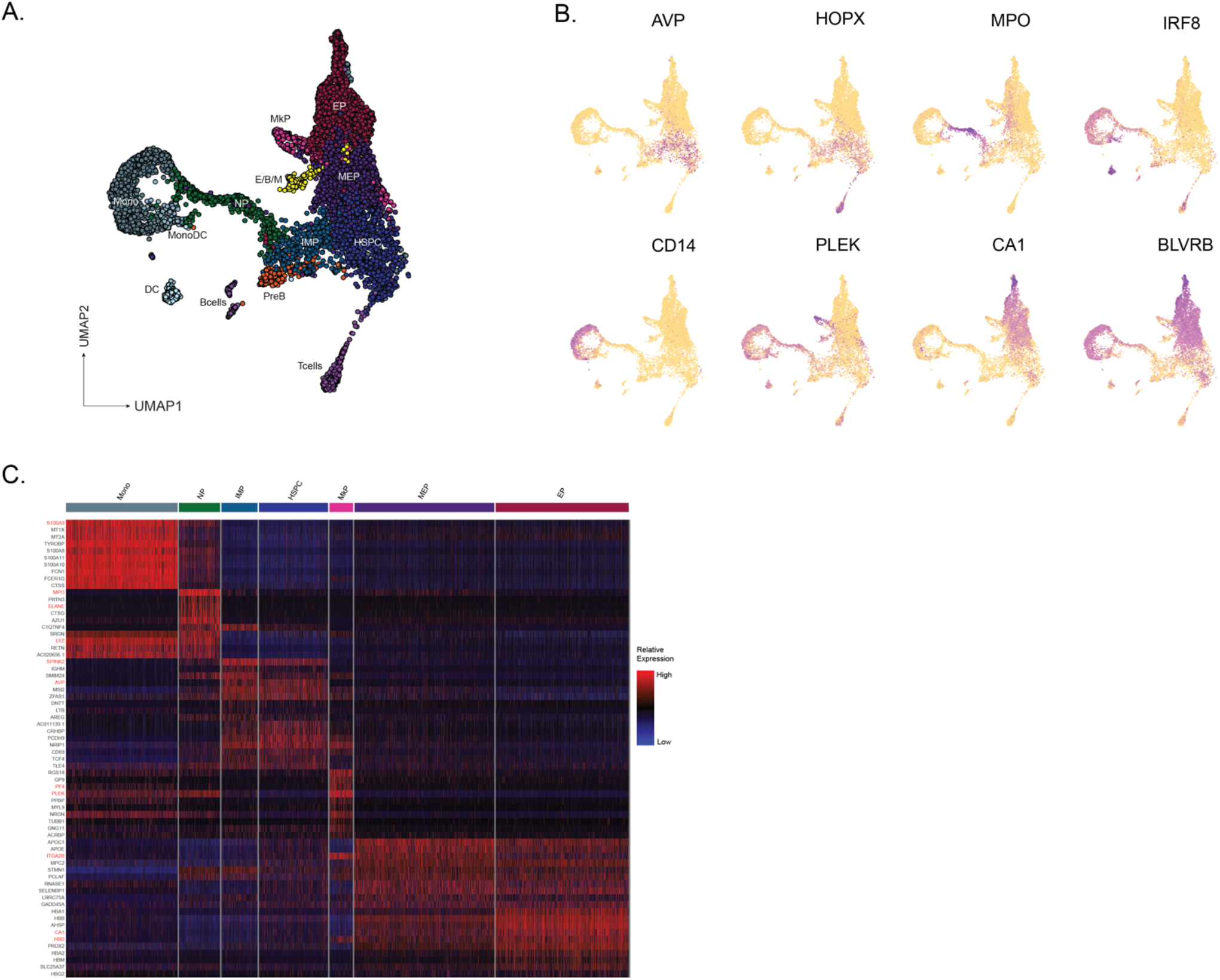
MDS validation clustering and cell-type assignment. **(A)** UMAP of CD34+ sorted cells (*n =* 8,879 cells) from samples MDS04-06 with *SF3B1* K700E mutations (*n =* 3), overlaid with cluster cell-type assignments. HSPC, hematopoietic stem progenitor cells; IMP, immature myeloid progenitors; MkP, megakaryocytic progenitors; MEP, megakaryocytic-erythroid progenitors; EP, erythroid progenitors; NP, neutrophil progenitors; E/B/M, eosinophil/basophil/mast progenitor cells; T/B cells; Mono, monocyte; DC, dendritic cells; Pre-B, precursors B cells; Mono DC, monocyte/dendritic cell progenitors. **(B)** Expression of lineage-specific genes from Velten et al.^37^ scored and projected onto the UMAP representation of cells from MDS04-06. **(C)** Heatmap of top 10 differentially expressed genes for each progenitor subset for MDS04-06.

**Extended Data Figure 4.**
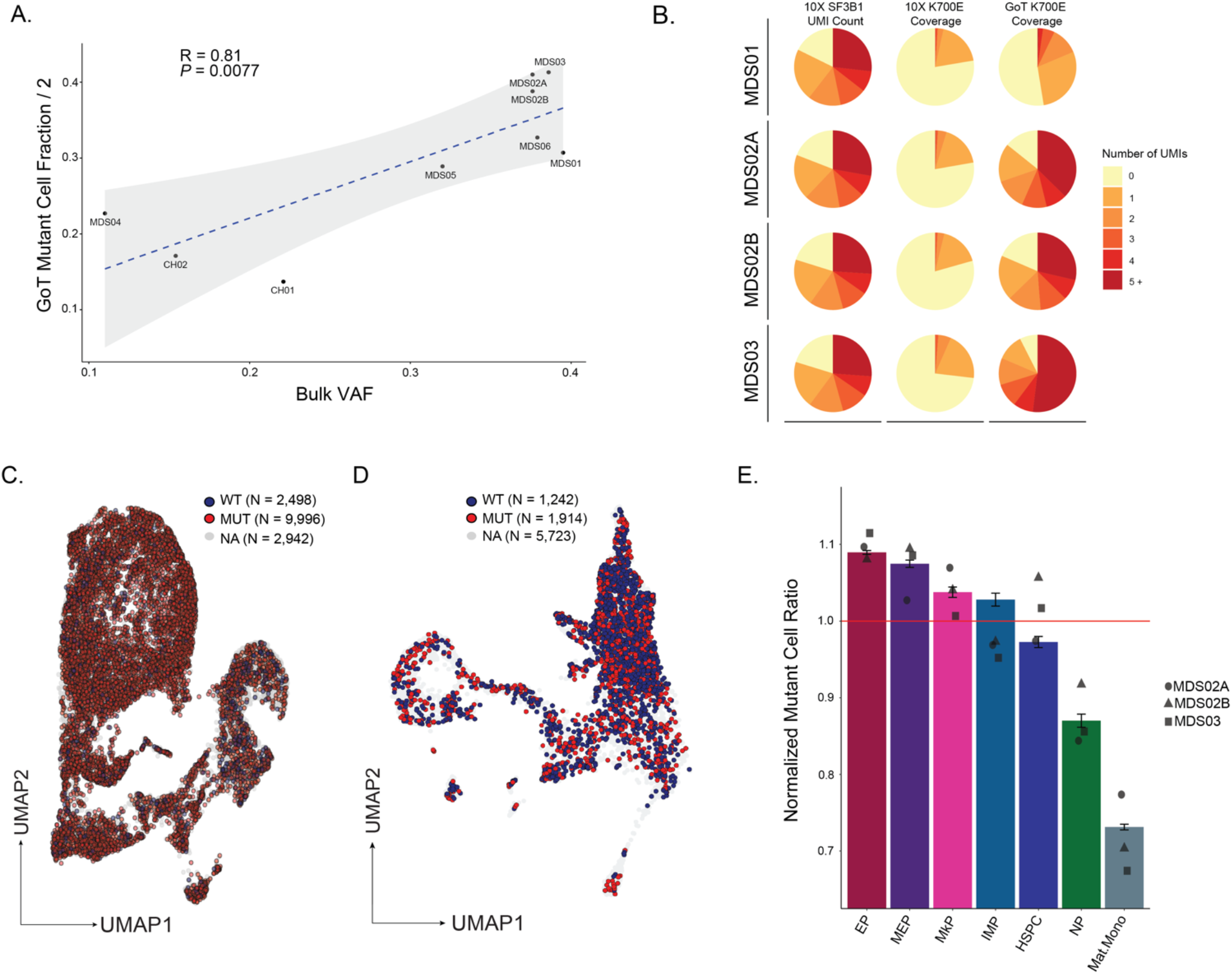
GoT statistics and analyses. **(A)** *SF3B1* K700E (7) and K666N (1) mutant cell fractions determined by GoT in single cells versus *SF3B1* K700E and K666N mutation variant allele frequencies (VAF) determined in bulk sequencing of matched unsorted bone marrow mononuclear cells (MDS) or matched unsorted stem cell product (CH). **(B)** Fraction of cells in MDS01-03 by number of *SF3B1* UMIs in standard 10x Genomics data without genotyping information (left), *SF3B1* UMIs with K700E locus coverage in standard 10x data (middle), and *SF3B1* UMIs with K700E locus coverage in GoT amplicon library (right). **(C)** UMAP of progenitor cells from MDS01-03 overlaid with genotyping data. WT, cells with genotype data without *SF3B1* mutation; MUT, cells with genotype data with *SF3B1* mutation; NA, unassignable cells with no genotype data. **(D)** UMAP of progenitor cells from MDS04-06 overlaid with genotyping data. WT, cells with genotype data without *SF3B1* mutation; MUT, cells with genotype data with *SF3B1* mutation; NA, unassignable cells with no genotype data. **(E)** Normalized ratio of *SF3B1*^*mut*^ cells in progenitor subsets with at least 300 genotyped cells. Bars show aggregate analysis of samples MDS01-03 with mean +/-s.e.m. of 100 downsampling iterations to 1 genotyping UMI per cell. Points represent the mean of *n* = 100 downsampling iteration per sample.

**Extended Data Figure 5.**
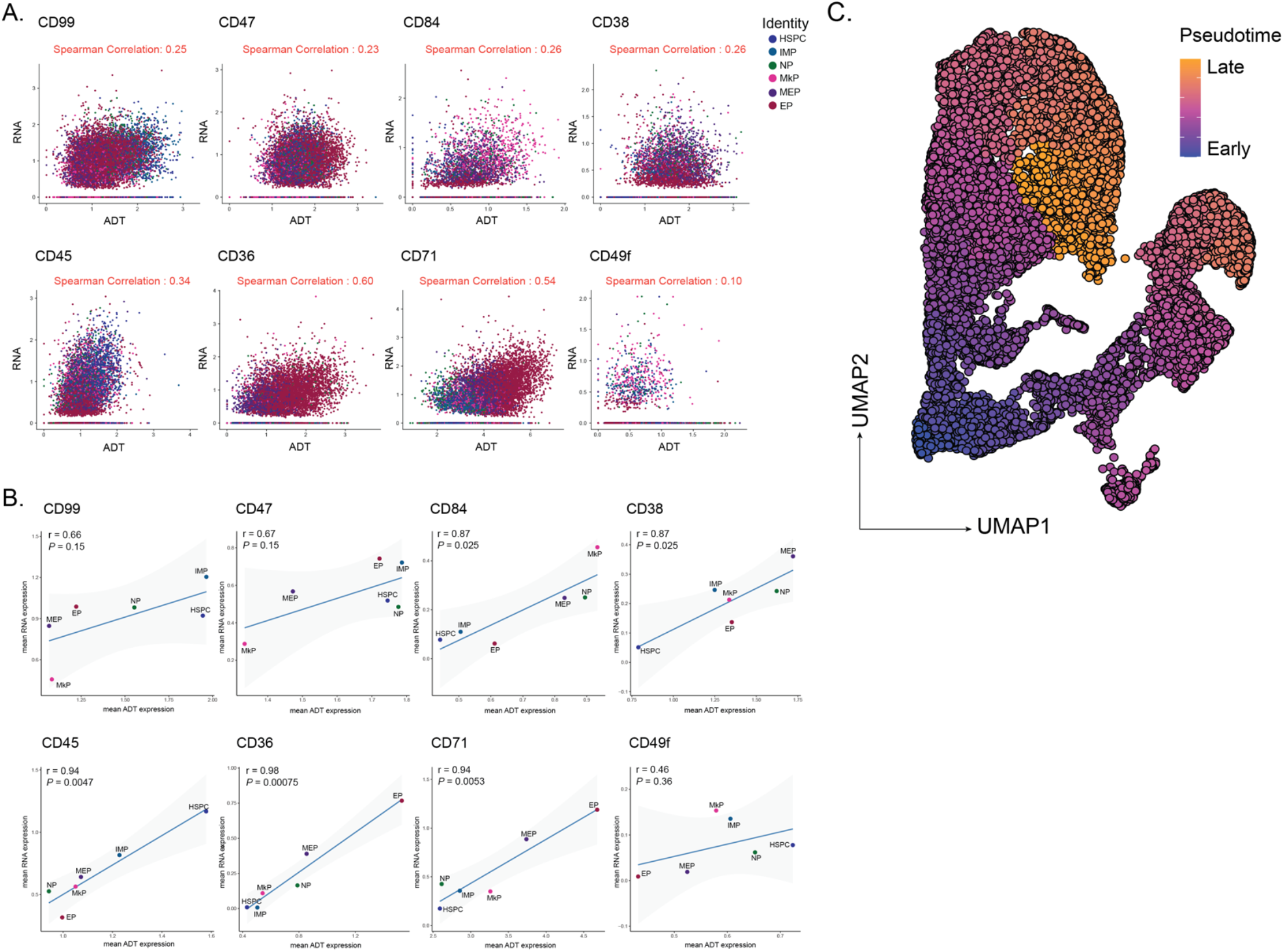
MDS CITE-seq and pseudotime. **(A)** Comparison of the single-cell expression of markers captured in both CITE-seq (x-axes) and RNA-seq (y-axes) libraries. Correlation coefficient r calculated using Spearman’s correlation. Cells are colored by each progenitor subset. **(B)** Comparison of the mean expression per progenitor subset of markers captured in both CITE-seq (x-axes) and RNA-seq (y-axes) libraries. Correlation coefficient r calculated using Spearman’s Correlation. *P*-values derived from Student’s t-distribution. **(C)** UMAP of progenitor cells from MDS01-03 samples overlaid with pseudotemporal ordering.

**Extended Data Figure 6.**
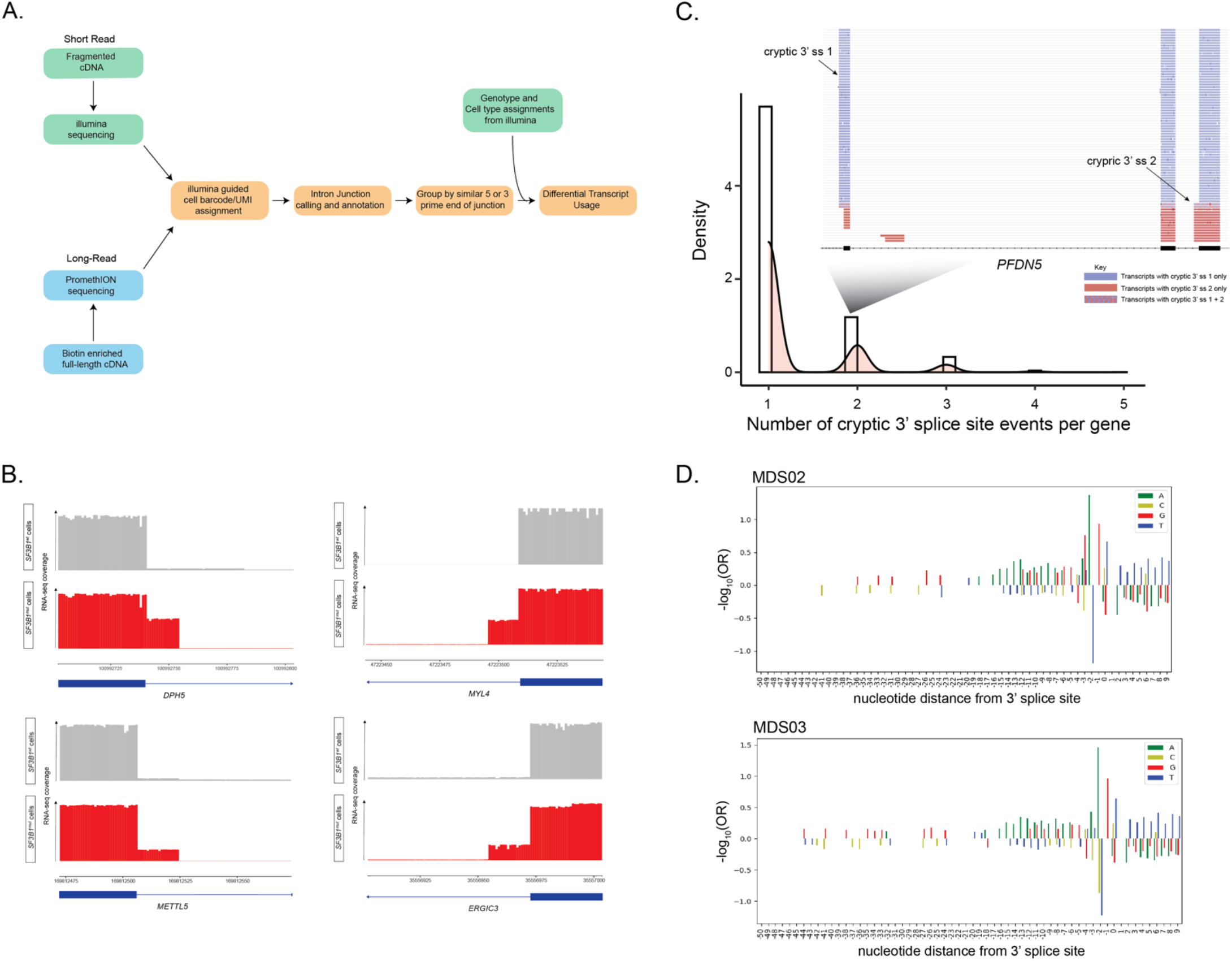
Long-read splicing and motif analysis. **(A)** Long-read sequencing processing and splicing analysis pipeline. **(B)** Comparison of the usage of various alternative 3’ splice sites found in our MDS *SF3B1*^*mut*^ cells vs. a CD34+ sample with no *SF3B1* mutation. **(C)** Bar plot of the number of cryptic 3’ splice sites identified per gene in MDS. *Inset*: Gene example, *PFDN5*, with 2 unique cryptic 3’ splice sites, showing the transcripts that have usage of either site. **(D)** Nucleotide enrichment (measured as log-odds ratio) across the 3’ splice site region comparing cryptic vs. canonical sites in MDS02-03 samples.

**Extended Data Figure 7.**
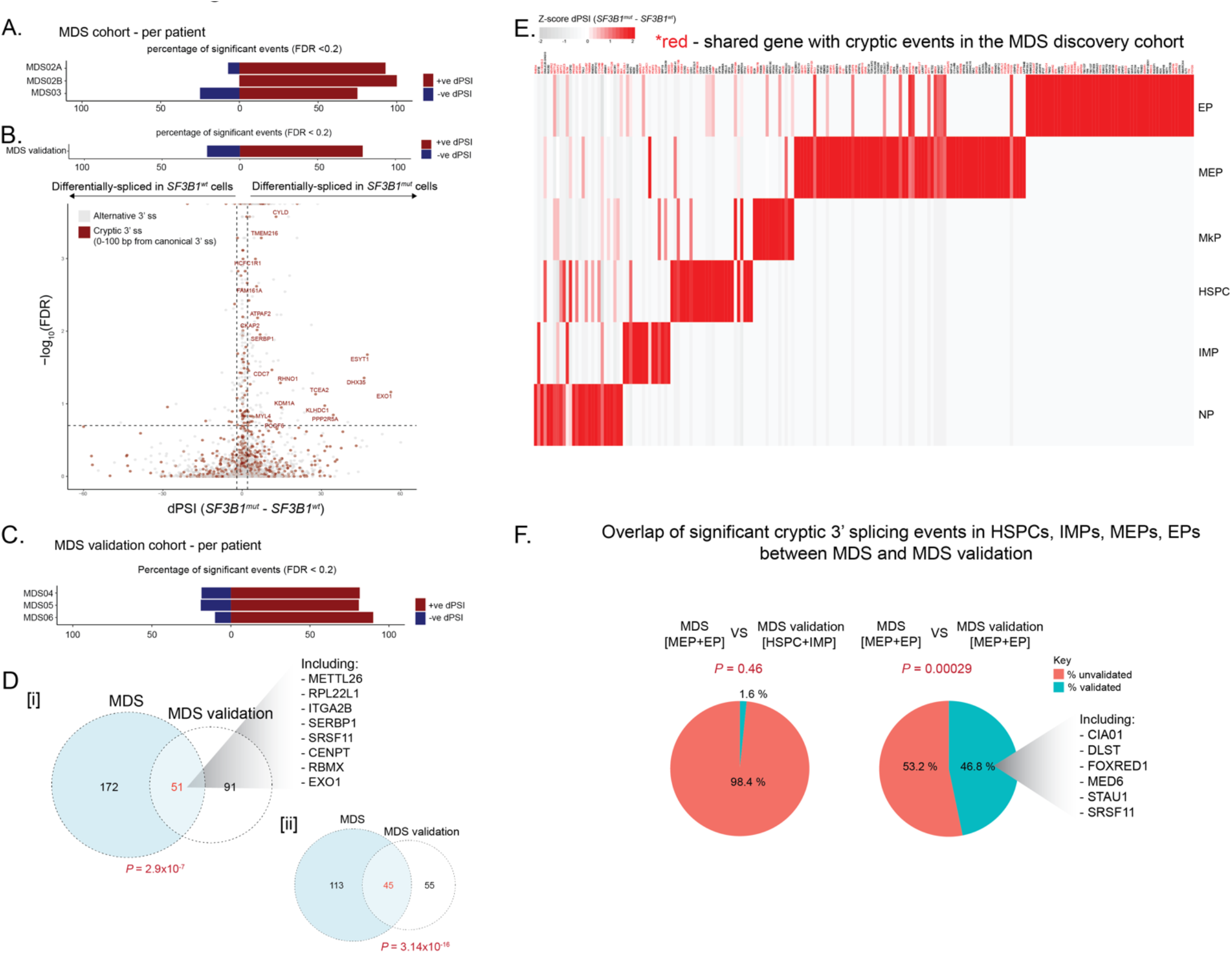
MDS vs. MDS validation cohort splicing comparison. **(A)** Bars showing the percentage of genes differentially spliced in *SF3B1*^*mut*^ and *SF3B1*^*wt*^ cells with BH-FDR adjusted *P*-value < 0.2 in each sample of the MDS cohort (MDS02(A/B)-03). **(B)** Differential splicing analysis between *SF3B1*^*mut*^ and *SF3B1*^*wt*^ cells across the aggregate of the MDS validation cohort (MDS04-06). Junctions with an absolute delta percent spliced-in (dPSI) > 2 and BH-FDR adjusted *P*-value < 0.2 were defined as differentially spliced. Bars (top) showing the percentage of genes differentially spliced in *SF3B1*^*mut*^ and *SF3B1*^*wt*^ cells of MDS validation cohort. **(C)** Bars showing the percentage of genes differentially spliced in *SF3B1*^*mut*^ and *SF3B1*^*wt*^ cells with BH-FDR adjusted *P*-value < 0.2 in each sample of the MDS validation cohort (MDS04-06). **(D)** Venn Diagram for the overlap of differentially spliced genes used more highly in *SF3B1*^*mut*^ cells (P-values < 0.05, dPSI >0) from the bulk comparison of *SF3B1*^*mut*^ vs. *SF3B1*^*wt*^ cells in the MDS and MDS validation cohorts [i]. Increasing the read coverage threshold for the differentially spliced genes showed a more significant overlap between cohorts [ii]. *P*-values for the overlap from Fisher’s exact test. **(E)** Heatmap of dPSI values between *SF3B1*^*mut*^ and *SF3B1*^*wt*^ cells for cryptic 3’ splicing events identified in the main progenitor subsets across MDS validation samples. Rows (z-score normalized) correspond to cryptic 3’ junctions found to be differentially spliced in at least one cell-type, with *P*-value <= 0.05 and dPSI >= 2. Columns correspond to cell-type. Genes with an *SF3B1*^*mut*^ associated cryptic 3’ splice site found in the MDS cohort highlighted (Red). **(F)** Pie chart showing the percent overlap of cryptically 3’ spliced genes unique to MEPs and EPs in the primary MDS cohort that are also cryptically 3’ spliced and unique to earlier progenitor cells (HSPCs and IMPs) in the MDS validation cohort (left) as well as the percent overlap with genes cryptically 3’ spliced and unique to the MEPs and EPs in the MDS validation cohort (right). *P*-value for the overlap from Fisher’s exact test.

**Extended Data Figure 8.**
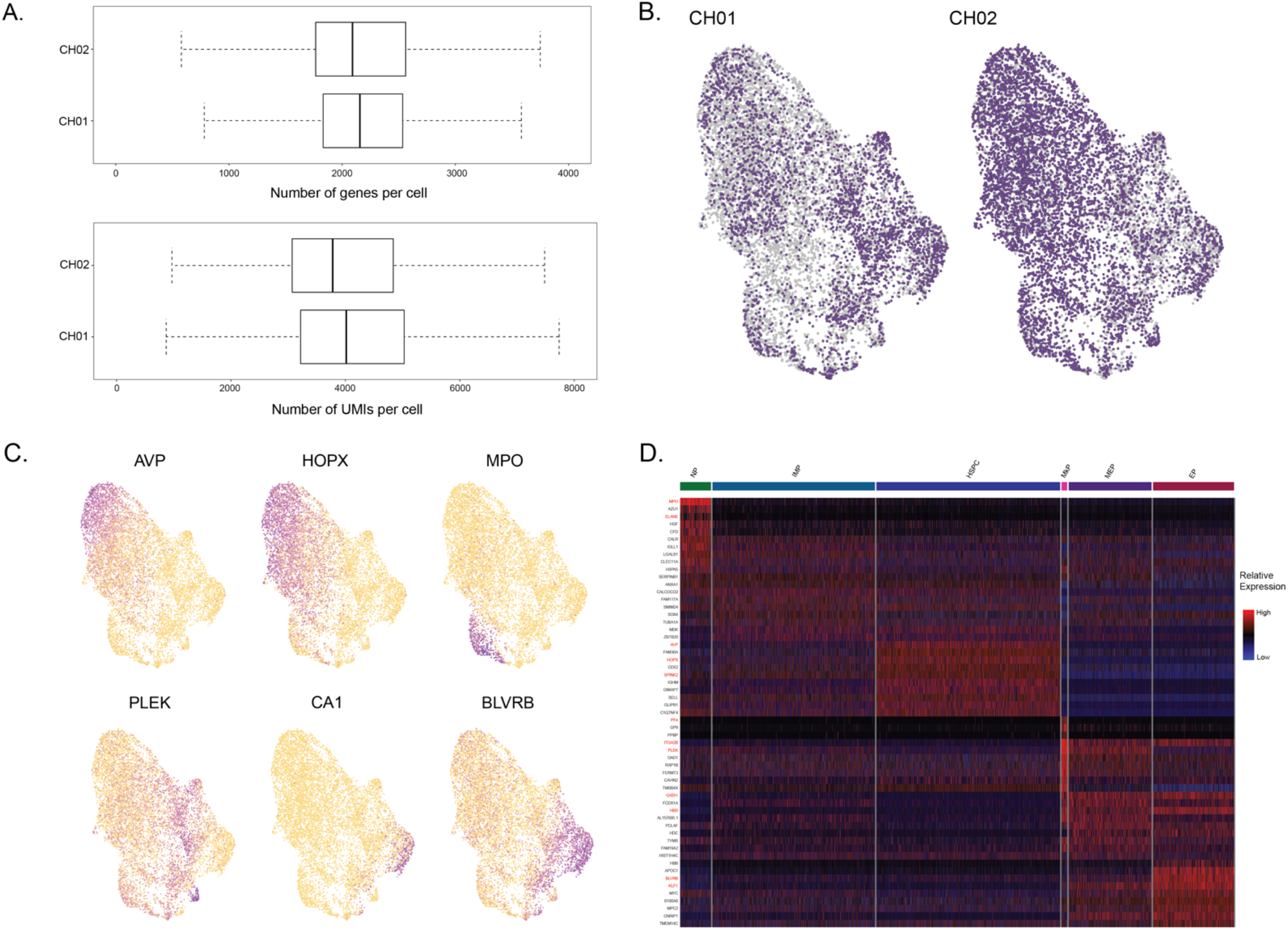
CH cohort QC and integration. **(A)** Number of genes per cell (top) and number of UMIs per cell (bottom) in CD34+ sorted hematopoietic progenitors from samples CH01-02 after QC filters, shown by each patient sample. **(B)** UMAP of CD34+ sorted progenitor cells for each individual sample of CH01-02 after integration using the Seurat package. **(C)** Expression of lineage-specific genes from Velten et al.^37^ scored and projected onto the UMAP representation of cells from CH01-02. **(D)** Heatmap of top 10 differentially expressed genes for each progenitor subset for CH01-02.

**Extended Data Figure 9.**
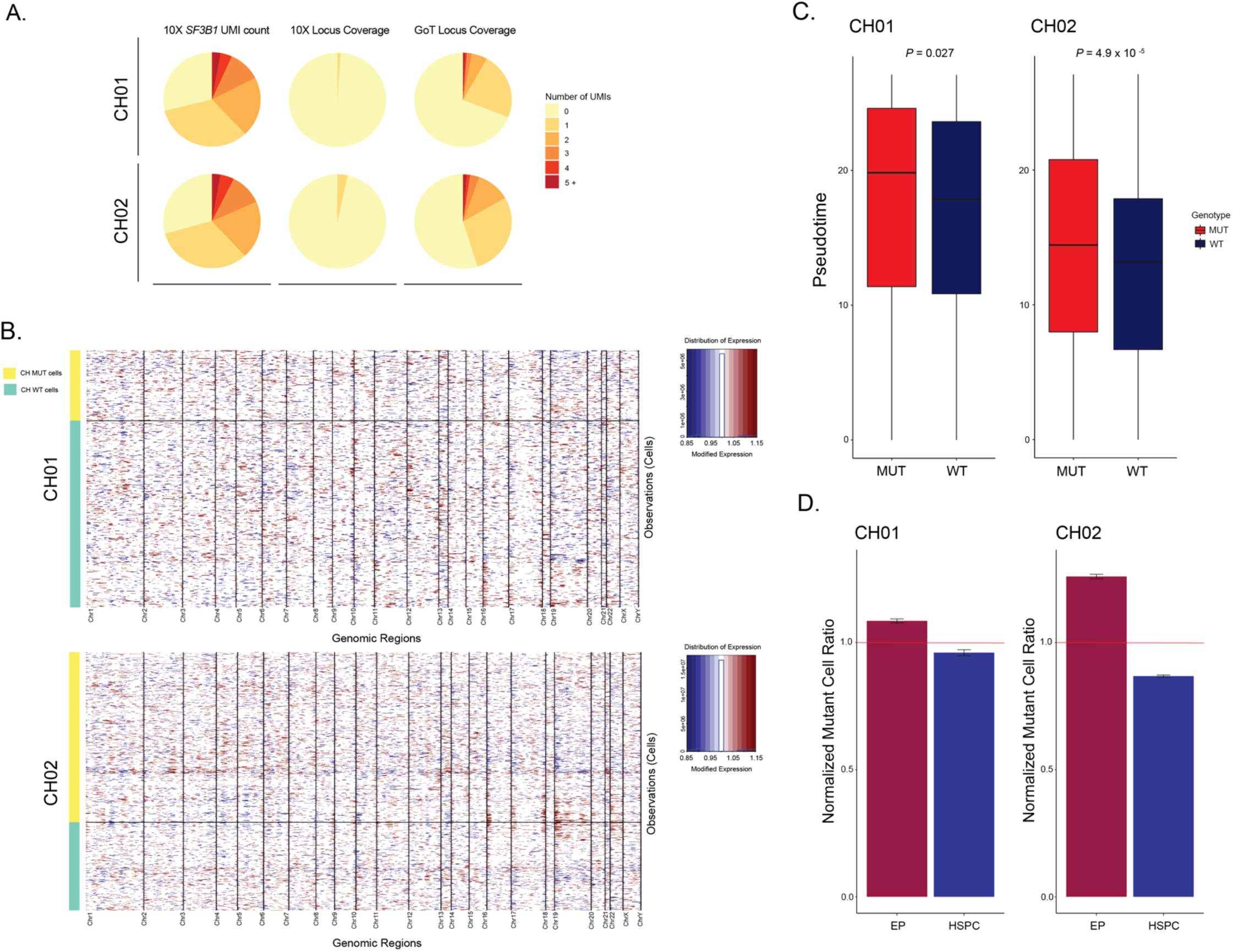
CH GoT statistics and analyses. **(A)** Fraction of cells in CH01-02 by number of *SF3B1* UMIs in standard 10x Genomics data without genotyping information (left), *SF3B1* UMIs with K666N (CH01) or K700E (CH02) locus coverage in standard 10x data (middle), and *SF3B1* UMIs with K666N (CH01) or K700E (CH02) locus coverage in GoT amplicon library (right). **(B)** Normalized ratio of *SF3B1*^*mut*^ cells in HSPC and EP cells for CH01 and CH02. Bars show the mean of *n =* 100 downsampling iterations to 1 genotyping UMI per cell. **(C)** Per sample heatmap of relative expression of genes ordered by chromosome/chromosomal position following copy number variation analysis using the InferCNV package (see Methods). Cells (y-axis) are stratified by *SF3B1* genotype status. **(D)** Pseudotime in *SF3B1*^*mut*^ vs. *SF3B1*^*wt*^ cells per CH sample. *P*-value for comparison of means from Wilcoxon rank sum test.

**Extended Data Figure 10.**
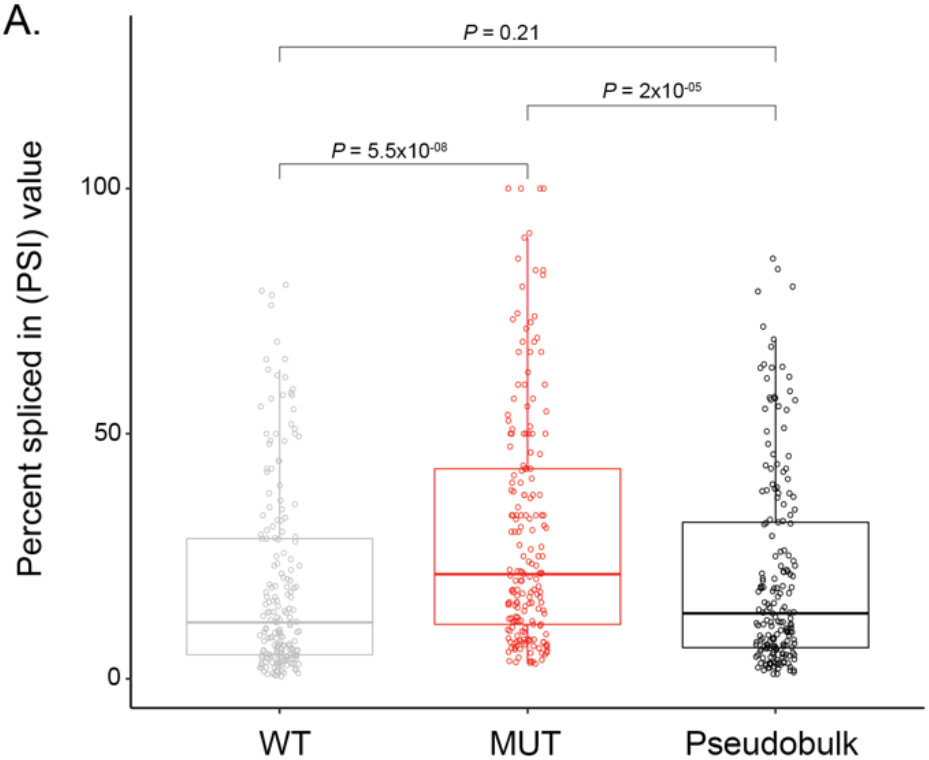
CH cryptic signal, WT/MUT/Pseudobulk comparison. **(A)** Comparison of the PSI values of identified cryptic junctions in WT cells only (gray) vs. MUT cells only (red) vs. all cells in pseudobulk in CH01-02. *P*-values for comparison of means from Wilcoxon rank sum test.

